# Experience-dependent place-cell referencing in hippocampal area CA1

**DOI:** 10.1101/2023.11.23.568469

**Authors:** Fish Kunxun Qian, Yiding Li, Jeffrey C. Magee

## Abstract

CA1 hippocampal place cells (PCs) are known for using both self-centric (egocentric) and world-centric (allocentric) reference frames to support a cognitive map^1,2^. The mechanism of PC referencing and the role of experience in this process, however, remain poorly understood^3–5^. Here we longitudinally recorded the activity of CA1 PCs while mice performed a spatial learning task. In a familiar environment, the CA1 representation consisted of PCs that were referenced to either spatial locations (allocentric PCs) or mouse running (egocentric PCs) in approximately equal proportions. In a novel environment, however, the CA1 representation became predominately egocentrically referenced. Notably, individual allocentric PCs in a familiar environment adaptively switched reference frames to become egocentric in a novel environment. In addition, intracellular membrane potential recordings revealed that individual CA1 neurons simultaneously received both ego- and allo-centric synaptic inputs, and the ratio of these two input streams correlated with the level of individual PC referencing. Furthermore, behavioral timescale synaptic plasticity^6,7^ (BTSP) was an active participant in shaping PC referencing through the rapid adjustment of synaptic weights on many PCs. Together, these results suggest that experience-dependent adjustment of synaptic input shapes ego and allocentric PC referencing to support a flexible cognitive map in CA1.

## Introduction

Goal-directed spatial navigation is an essential brain function and it has been observed that animals use two distinct sets of reference frames during navigation. First, movement produces selfmotion cues, the integration of which (i.e., path integration) allows the animal to internally register its position using the self-centered (egocentric) reference frame^2,8–11^. However, egocentric spatial coding using path integration alone is prone to accumulated drift errors over time^2,8–11^. Alternatively, an animal can use external cues and landmarks to provide world-centered (allocentric) reference frames^1,2,11^ that can correct path integration errors and more precisely register the animal’s position in an environment^1,2,10–12^. Thus, egocentric and allocentric reference frames are distinct yet complementary for registering spatial location. However, it remains unclear how the brain integrates different reference frames during navigation.

The hippocampus is a brain region critical for spatial navigation and episodic memory^13^. Place cells (PCs) in the hippocampus robustly increase their firing rates when an animal enters a specific place (place field; PF) in the environment^14^. Three major lines of evidence support the idea that PCs in rodents primarily rely on allocentric reference frames. First, PCs recorded from freely behaving animals are preferentially controlled by external cues which collectively provide an allocentric reference frame^8,15–18^. Second, PCs are insensitive to the animal’s specific route into the PF location (omni-directional) when an animal is randomly foraging in an open field without a goal^8,19^. Third, the medial entorhinal cortex provides spatially tuned input to the hippocampus that is thought to supply the basis for allocentric PCs^20,21^. Therefore, the collective activity of PCs (i.e., PC representation) is believed to serve as a neural substrate for an allocentric cognitive map of space in the brain.

On the other hand, evidence supporting an important role for egocentric reference frames in registering hippocampal PC representations is accumulating^3,22,23^. First, the PC representation remains largely intact when external visual cues are eliminated, suggesting that egocentric frames of reference are sufficient to maintain a learned spatial map in the hippocampus^8,17,24,25^. Moreover, PCs become directional^26,27^ and their PFs reorganize after a goal location has been introduced within an environment^27–31^. In addition, PCs can also primarily encode the distance an animal travels from the previously occupied position in an egocentric manner^32–34^ through the process of path-integration^2,10–12^. In this case, these egocentric PCs will fire at a particular distance that the animal has traveled from its start position. Finally, the lateral entorhinal cortex provides robust egocentric spatial information to the hippocampus that may supply the basis for egocentric PCs^35^. Therefore, the hippocampus might be well suited for integrating both egocentric and allocentric reference frames to produce a flexible cognitive map of space. To examine this issue, we sought to specifically determine the reference frame of individual PCs in a CA1 representation of the environment, the degree to which this referencing is plastic and its synaptic mechanisms.

## Results

To determine the contribution of allocentric and egocentric reference frames to PC activity requires independent experimental manipulations of the different reference frames^3,4^. Here we recorded from adult mice engaging in a hippocampal-dependent goal-directed spatial learning task on a linear treadmill^36,37^. The task requires head-fixed mice to run for a sucrose water reward (delivered at 50 cm) on a 180-cm-long track (belt A), uniformly covered with distinct tactile cues to aid navigation (Fig. 1a). After training, mice typically developed anticipatory slowing and licking before the reward delivery site, indicating that the location of the reward was successfully learned (Fig. 1b, Pre: before reward switch). We performed *in vivo* two-photon calcium imaging^38^ to optically record populations of PC activity from CA1 pyramidal neurons that transgenically expressed GCaMP6f^39^ (Fig. 1a, middle and right). We observed large numbers of PCs per animal (90.2±14.2 PCs/animal, n = 6 mice) that exhibited reliable trial-by-trial fluorescence increases at selective locations^1,31,37,40,41^ (Fig. 1c, Pre). Consistent with previous studies, these PCs tiled the entire track with an elevated density found around the reward location (i.e., over-representation; Extended Data Fig. 1a)^30,31,37,42–45^.

**Fig. 1.**
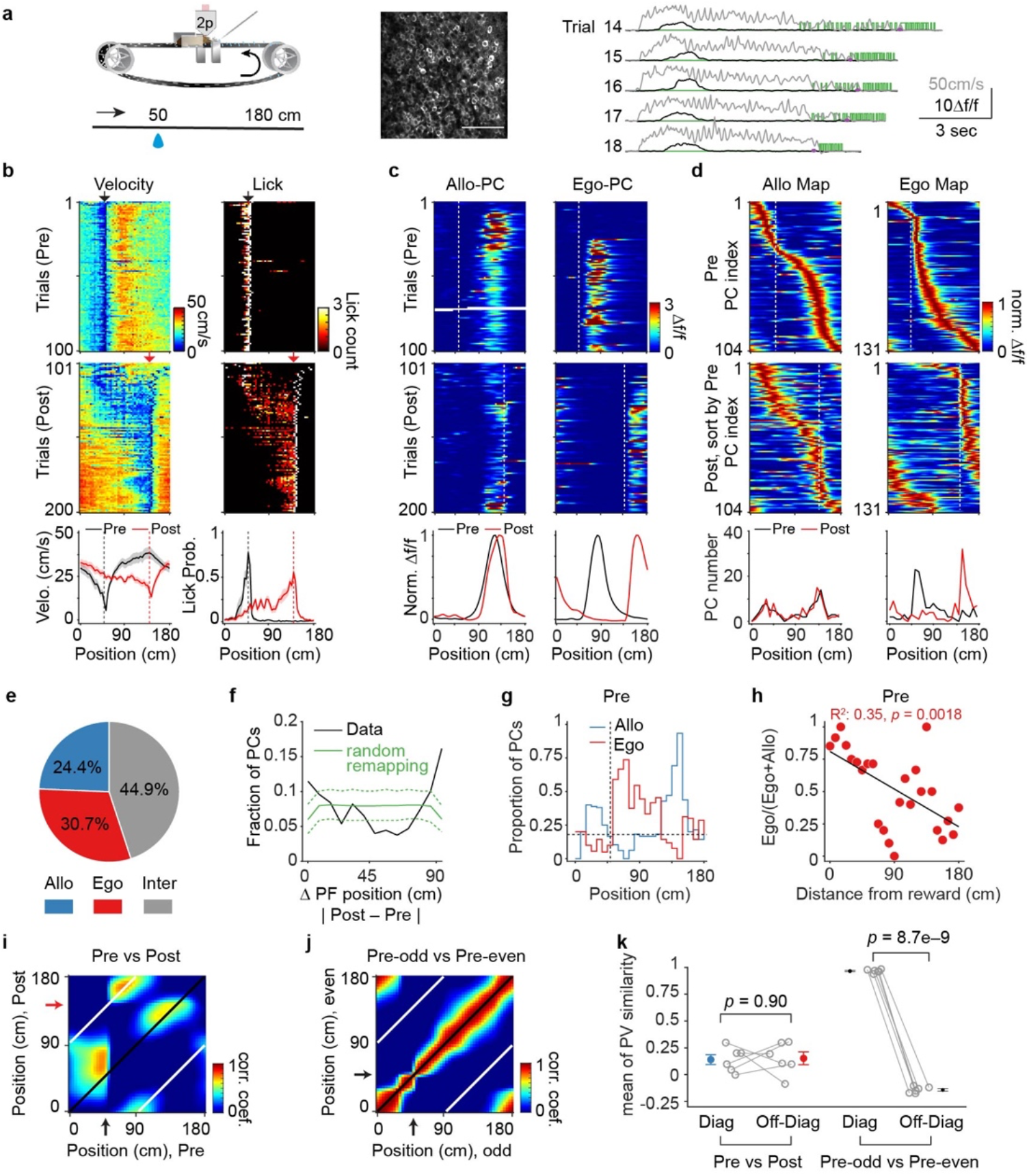
Balanced egocentric and allocentric spatial coding in a familiar environment. **a**, Left: mouse running on a linear treadmill to navigate to water reward at a fixed location. Middle: example time-averaged image of field of view under two-photon microscope showing GCamp6f-expressing pyramidal neurons in dorsal CA1. Scale bar: 100 μm; Right: example traces from 5 trials illustrating Δf/f from a PC (black), running velocity (grey), and licks (green). Purple dots mark the reward location. **b,** Top and middle: running (left) and licking (right) behavior of a representative animal on a familiar belt. Pre: before the reward switch. Post: after the reward switch. Arrows mark the reward locations during Pre (black, at 50 cm) and Post (red, at 140 cm) sessions. Bottom: mean velocity (left) and licking probability (right) of all mice (n = 6) during Pre and Post, respectively. Shading represents SEM. Reward locations are marked by vertical dashed line. **c**, Example of one allocentric PC (left) and one egocentric PC (right) illustrating Δf/f in space across laps during Pre (top) and Post (middle). Vertical dashed white lines mark the reward locations. Bottom: peak-normalized mean Δf/f across space for Pre and Post, respectively. **d,** PC activity on a familiar belt. Top: peak-normalized mean Δf/f of all allocentric PCs (left) and egocentric PCs (right) in the Pre condition. PCs are sorted by the peak location of mean Δf/f. Middle: peak-normalized mean Δf/f of the same allocentric and egocentric PCs in Post, sorted by Pre location. Vertical dashed white lines mark the reward locations. Bottom: number of PCs across space during Pre and Post for allocentric PCs (left) and egocentric PCs (right). **e**, The fraction of different PC activity profiles (n = 427 PCs from 6 mice). **f,** Fraction of PCs as a function of distance of PF shift for measured and simulated data. Green dashed lines mark the top and bottom 5^th^ percentile of the simulated values using a random remapping model. **g,** Proportion of allocentric and egocentric PCs, for all PCs, across space in Pre. Vertical dashed line marks the reward location. Horizontal dashed line marks the expected fraction (∼0.167) if the remapping was random. **h,** Ratio of egocentric PCs over the sum of egocentric and allocentric PCs across distance from reward site in Pre. Black line is a linear fit. **i.** Correlation matrix using Pearson’s correlation coefficient between averaged population vectors (PVs) at all locations from Pre and Post. Black line marks the diagonal, white lines mark the off-diagonal. **j,** Same analysis as **i**, but between PVs averaged from odd trials and even trials in Pre. **k,** Mean of the Pearson’s correlation coefficient between PVs along the diagonal and off-diagonal axis for all animals. (Pre vs Post, diagonal: 0.14±0.046, off-diagonal: 0.15±0.060. Paired two-sample *t*-test, *p* = 0.90) (Pre-odd vs Pre-even, diagonal: 0.97±0.0065, off-diagonal: –0.14±0.012. Paired two-sample *t*-test, *p* = 8.7e–9)

### Balanced allocentric and egocentric spatial coding in a familiar environment

In our spatial learning task, distinct tactile cues on the belt potentially provide allocentric reference points^44,46^, while a mouse’s stereotypical running pattern provides an egocentric reference frame where integration of self-motion over time is, theoretically, sufficient for the animal to update its position from the beginning of the running^10–12^. To investigate whether the CA1 PC representation references to egocentric or allocentric reference frames in our task, we switched the reward from 50 cm to 140 cm (half of the belt length) after the first 100 trials to alter the relationship between the running pattern of the animal and the spatial cues. This manipulation dissociates the animal’s egocentric and allocentric reference frames allowing us to study their respective influences on each PC’s activity^32,47–50^. As the mice adapted their anticipatory slowing and licking to the new reward location. As the mice adapted their anticipatory slowing and licking to the new reward location (Fig. 1b, Post: after reward switch), we observed dramatic PC remapping in CA1^15,31,51,52^; such that PCs changed their PF locations, preexisting PCs disappeared, and new PCs emerged (Extended Data Fig. 2b–c). Of those PCs with a reliable PF both before and after reward switch (65 % of all PCs, n = 6 mice, Extended Data Fig. 2c), we identified 24.4 % as allocentric PCs that maintained a stable PF relative to the spatial cues after the reward switch (PFs shifted <15 cm; Fig. 1c, left; 1d, left; 1e, blue; see Extended Data Fig. 2a for criterion), 30.7 % as egocentric PCs that shifted their PFs in space but maintained the same relation to the beginning of running on each lap (PFs shifted between 75–90 cm; Fig. 1c, right; 1d, right; 1e, red), and a remaining 44.9 % that were intermediate because their PFs shifted between 16 and 74 cm (Fig. 1e, Inter, grey; Extended Data Fig. 3a–b). The population distribution of PF displacement displayed two peaks toward 0 and 90 cm, indicating that PC remapping in CA1 was not random, but instead comprised of PCs that were primarily either allocentric or egocentric referenced (Fig. 1f).

Previous studies suggested that the egocentric spatial coding degrades progressively as the animal traverses away from its original reference point, while allocentric sensory cues, which are fixed in external space, can correct errors accumulated during navigation^10–12^. We therefore examined the spatial distribution of egocentric and allocentric PCs and found that egocentric PCs preferentially clustered near the onset of running (peak at 60 cm, that is, 10 cm after the reward site; Fig. 1d, right; 1g, red), while allocentric PCs were predominate over the rest of the track (peak at 144 cm and 36 cm; Fig. 1d, left; 1g, blue). Notably, more than half of the egocentric PCs had PFs located 30 cm distant in space (median: 32 cm; Fig. 1d, right) and all were many seconds in time from the reward site (time between reward delivery and the start of running: 6.8±0.19 sec, n = 6 mice, first 100 laps before reward switch), indicating that they did not simply represent appetitive reward responses^31,53^. At further distances from the reward site, the proportion of egocentric versus allocentric PCs decreased (Fig. 1h).

To examine how PC remapping influenced ensemble spatial coding before and after reward switch, we performed a population vector analysis^54^. In the correlation matrix formed from before and after reward switch activity, the single region of high correlation along the off-diagonal (white line, Fig. 1i) corresponded to egocentric PCs that primarily clustered at 60 cm (∼10 cm after the reward site; Fig. 1i; 1d, right), suggesting that the PC representation immediately after the running onset is primarily egocentric^32,34^. In contrast, two regions of high correlation along the diagonal (black line, Fig. 1i) corresponded to the allocentric PCs that clustered at ∼30 cm and ∼140 cm (Fig. 1i; 1d, left; 1g, blue), reflecting the predominance of allocentric PCs in coding these positions relative to the spatial cues on the belt. This can be compared with the trial-by-trial correlations which show a single high correlation value around the diagonal (Fig. 1j–k). The mean of the population vector correlation along the diagonal was nearly identical to that of the off-diagonal, suggesting that allocentric and egocentric reference frames have similar contributions to the ensemble spatial coding of the familiar environment (Fig. 1k, left). These results demonstrate that in a familiar environment both allocentric and egocentric reference frames exist supporting a coherent spatial representation in CA1.

### Experience alters the referencing of the CA1 PC representation

Next, we sought to examine how experience influences the balance of allocentric and egocentric spatial representation in CA1^55^. We trained a separate cohort of mice with fixed reward at 50 cm (n = 6 mice) on one belt (belt A, familiar), then exposed them to another belt covered with a set of unfamiliar tactile cues (belt B, novel). On the first day of exposure to the novel belt, mice quickly adapted their behavior (Fig. 2a, Pre). The PC representation of the novel belt also contained an over-representation around the reward site similar to that of a familiar belt (Extended Data Fig. 1b–c). PCs on a novel belt, however, had somewhat less spatial information and reduced trial-by-trial reliability (Extended Data Fig. 1d–e), consistent with previous reports^36,56–58^.

**Fig. 2.**
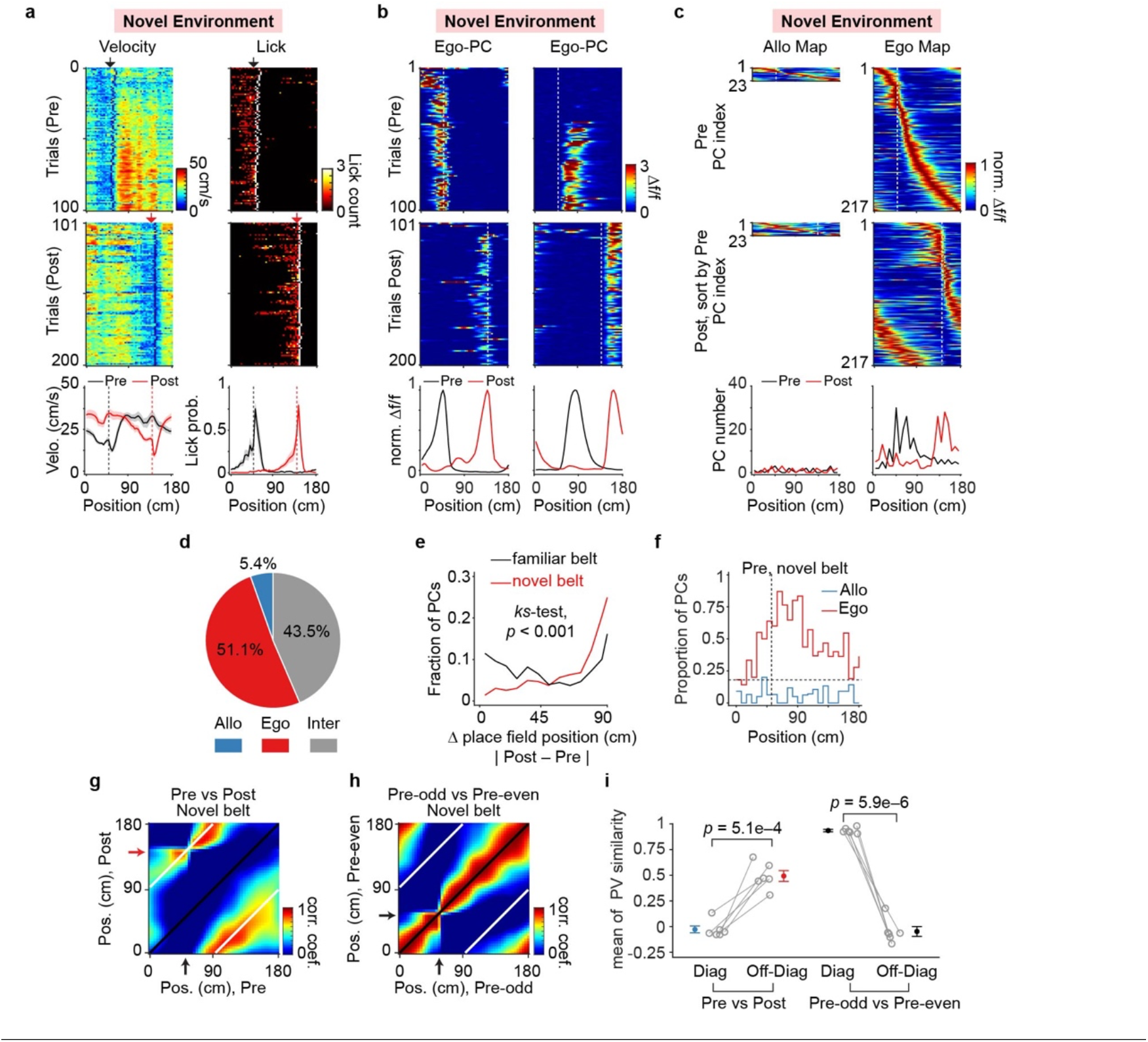
Predominant egocentric spatial coding in a novel environment. **a,** Top and middle: example running (left) and licking (right) behavior of an animal on a novel belt. Pre: before reward switch. Post: after reward switch. Arrows mark the reward locations during Pre (black, at 50 cm) and Post (red, at 140 cm). Bottom: mean velocity (left) and licking probability (right) of all mice (n = 6). Shading represents SEM. **b,** Example Δf/f activity in space across laps during Pre (top) and Post (middle) from two egocentric PCs on a familiar belt; one had a PF before and the other after the reward location. Vertical dashed white lines mark the reward locations. Bottom: peak-normalized mean Δf/f across space for Pre and Post, respectively. **c,** PC Δf/f activity on a novel belt. Top: peak-normalized mean Δf/f of all allocentric PCs (left) and egocentric PCs (right) in Pre. PCs are sorted by the peak location of mean Δf/f. Middle: peak-normalized mean Δf/f of the same allocentric and egocentric PCs in Post, sorted by Pre. Vertical dashed white lines mark the reward locations. Bottom: number of PCs across space during Pre and Post for allocentric and egocentric PCs, respectively. **d**, The fraction of different PC activity profiles (n = 425 PCs from 6 mice). **e,** Fraction of PCs as a function of distance of PF shift on the familiar (black, n = 6 mice) and novel (red, n = 6 mice) belt. (Kolmogorov-Smirnov test, *p* = 1.8e–17). **f,** Proportion of allocentric and egocentric PCs, over all PCs, across space. Vertical dashed line marks the reward location. Horizontal dashed line marks the expected fraction (∼0.167) if the remapping was random. **g,** Correlation matrix using Pearson’s correlation coefficient between PCs at all locations from Pre and Post at all locations on a novel belt. Black line marks the diagonal, white lines mark the off-diagonal. **h,** same analysis as **g**, but between PVs averaged from odd trials and even trials in Pre. **i,** Mean of the PV correlation coefficient along the diagonal and off-diagonal across animals on the novel belt (n = 6 mice). (Pre vs Post, diagonal: – 0.028±0.033, off-diagonal: 0.49±0.052. Paired two-sample *t*-test, *p* = 5.1e–4) (Pre-odd vs Pre-even, diagonal: 0.93±0.011, off-diagonal: –0.048±0.048. Paired two-sample *t*-test, *p* = 5.9e–6).

To understand how the manipulation of the allocentric reference frame (i.e., familiar to novel external cues) influences the balance of allocentric and egocentric PCs in the CA1, we next switched the reward location on the novel belt as previously described. Mice adapted their behavior as before (Fig. 2a, Post) but in stark contrast to the balanced proportions of allocentric and egocentric PCs on a familiar belt (Fig. 2e, black), the fraction of egocentric PCs on a novel belt was nearly ten times that of allocentric PCs (Fig. 2b–d; Extended Data Fig. 2d–e). Thus, PF displacement on a novel belt was heavily skewed toward 90 cm (Fig. 2e, red) and surprisingly, egocentric PCs tiled the entire novel belt (Fig. 2b; 2c, right; 2f, red). In addition, the fraction of allocentric PCs had a strong inverse correlation with the fraction of egocentric PCs across animals (Extended Data Fig. 3e), indicating that these two reference frames might compete with each other^32^. Moreover, population spatial coding was reliable on a novel belt before reward switch and exhibited egocentric dominance at all regions (Fig. 2g–i). Thus, under our experimental conditions, the PC representation in CA1 is primarily egocentric in a novel environment.

Importantly, the decreased proportion of allocentric PCs on a novel belt was not due to a lack of experience, as animals trained on the novel belt for another 4 consecutive days still did not develop balanced allocentric and egocentric referencing (n = 4 mice, Extended Data Fig. 4a). Also, the distinct allocentric and egocentric PC proportions on the familiar and novel belts were not due to the animals’ intrinsic preferences for different cues^59^ (n = 3 mice, Extended Data Fig. 4b). Together, these results demonstrate that experience alters the balance of egocentric and allocentric referencing present in the spatial representation of environments in CA1, and that exposure to a novel environment reshapes the representation to become primarily egocentric.

### Individual PCs change referencing

The above data suggest that individual PCs may change their referencing. To examine this we determined, as above, the PC referencing over multiple days under the two environmental conditions (day 1/day 2; familiar/familiar and familiar/novel). On the first day, we trained a set of mice on a familiar belt and switched the reward on this belt and categorized the referencing of the recorded PCs (familiar, day 1; Fig. 3a, left). On the next day (day 2, the mice were spilt into two separate groups, one that experienced a reward shift on the same familiar belt (Familiar/Familiar) and another that experienced the reward shift on a novel belt (Familiar/Novel). We recorded the same PCs on each belt and determined their referencing (day 2; Fig. 3a, right).

**Fig. 3.**
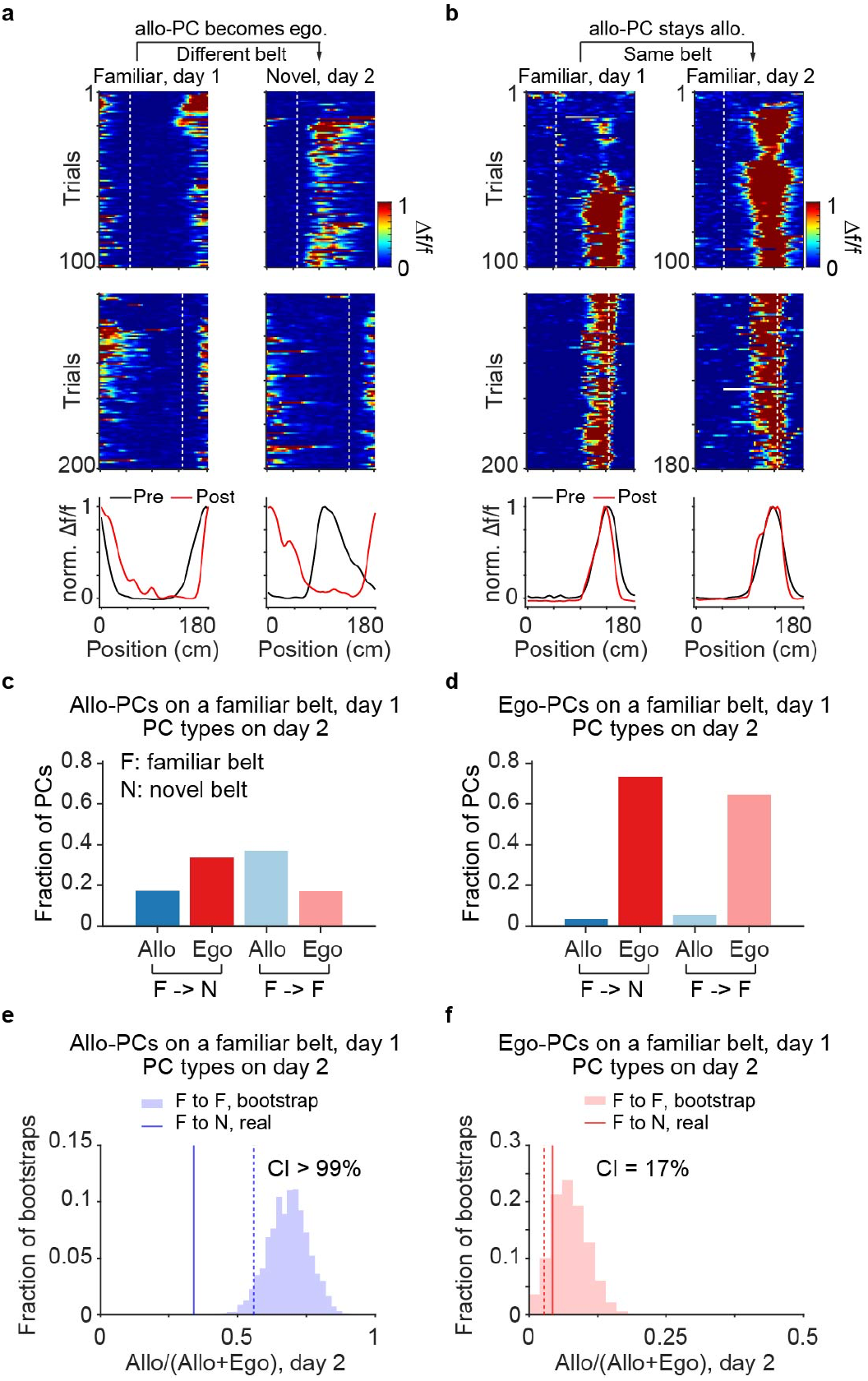
Individual PCs changed reference frames. **a,** Example of an allocentric PC switching to egocentric. The Δf/f activity across space for day 1 on a familiar belt (left) and day 2 on a novel belt (right) during Pre (top) and Post (middle) is shown. White dashed lines mark the locations of reward, 50 cm in Pre and 140 cm in Post. Bottom: peak-normalized Δf/f across space during Pre and Post for day 1 (familiar, left) and day 2 (novel, right). **b,** Same plots as in **a** showing a PC that remained allocentric on the same familiar belt for two consecutive days. **c,** Stability of allocentric PCs across 2 days. Left: fraction of allocentric PCs (day 1, familiar belt) that remained allocentric (dark blue) or switched to egocentric (dark red) on day 2 on a novel belt. Right: fraction of allocentric PCs (day 1, familiar belt) that remained allocentric (light blue) or became egocentric (light red) on day 2 on the same familiar belt. **d,** Same plot as **c**, showing stability of egocentric PCs across 2 days. **e,** Illustration of the stability of allocentric PCs across two days. The resampled distribution illustrates the fraction of bootstrapped values as a function of an Allo/(Allo+Ego) index when mice remained on the same familiar belt for two days (F to F: familiar to familiar). “Allo/(Allo+Ego)” index is calculated as the number of allocentric PCs that stayed allocentric on next day divided by the total number of allocentric PCs that stayed allocentric or became egocentric on next day. For each bootstrap, we sampled a fraction of the real data with replacement to calculate the Allo/(Allo+Ego) index using allocentric PCs on day 1. Dashed line marks 5^th^ percentile of the bootstrapped values during F to F. Solid line marks measured proportion when animals switched belt from familiar to novel (F to N: familiar to novel), indicating that significantly more allocentric PCs from day 1 became egocentric on day 2 on a novel belt (n = 6 mice) than animals that stayed on the same familiar belt for two days (n = 6 mice). CI: confidence interval. **f,** Same analysis as in **e**, but testing the stability of egocentric PCs across two days.

Of allocentric PCs on a familiar belt that remained PCs after the reward switch on a novel belt (familiar/novel; 64 % of all allocentric PCs, Extended Data Fig. 5a), 34 % became egocentric (Fig. 3a), twice as many as those that remained allocentric (17 % stayed allocentric; Fig. 3c, left). In contrast, when the same familiar belt was employed on both days, 37 % of allocentric PCs that remained PCs on the same familiar belt on the next day remained allocentric (familiar/familiar; 73 % of all allocentric PCs; Fig. 3b; Extended Data Fig. 5b), whereas 17 % became egocentric (Fig. 3c, right). Therefore, allocentric PCs remained relatively stable on the same belt across two days, with a small but noticeable proportion of reference frame switching (Fig. 3e; Extended Data Fig. 6). However, the introduction of a novel belt markedly increased the rate of reference frame switching from allocentric PCs to egocentric PCs (Fig. 3c; 3e, CI > 99 %; Extended Data Fig. 6). On the other hand, egocentric PCs on day 1 remained stable on the next day for both conditions (Fig. 3d–f; Extended Data Fig. 5c–d; 7a–b). These results demonstrate that experience flexibly shapes individual PCs to switch references between allocentric and egocentric reference frames.

### PCs receive both egocentric and allocentric referenced synaptic inputs

We next questioned what are the mechanisms controlling PC referencing? Given that modifications in the strength of CA3 synaptic input produce the membrane potential (V_m_) depolarizations that drive CA1 PF activity^6,60–63^, we turned to intracellular V_m_ recordings^61,64^ to examine how the synaptic input to PCs was affected by reward location changes. We performed whole-cell V_m_ recordings from CA1 pyramidal neurons of mice running on a familiar belt (Fig. 4a). In all recorded neurons intracellular currents were injected to induce new PFs as previously reported^6,61,65^. Consistent with previous intracellular recordings^61–64,66,67^, these PCs exhibited slow ramps of depolarization that drove PF firing at selective locations (V_m_ ramps; mean ramp amplitude, 8.89±0.64 mV, n = 26 cells; Extended Data Fig. 8b). We next switched the reward site as above and found that these PCs showed diverse PF remapping (6/26 were egocentric, 10/26 were allocentric, and 10/26 were intermediate; Extended Data Fig. 8a). After reward switch the location of V_m_ ramps was strongly correlated with PF locations (Extended Data Fig. 8c) and the distribution of PF displacement was consistent with that of PCs recorded via two-photon Ca^2+^ imaging (Extended Data Fig. 8d).

**Fig. 4.**
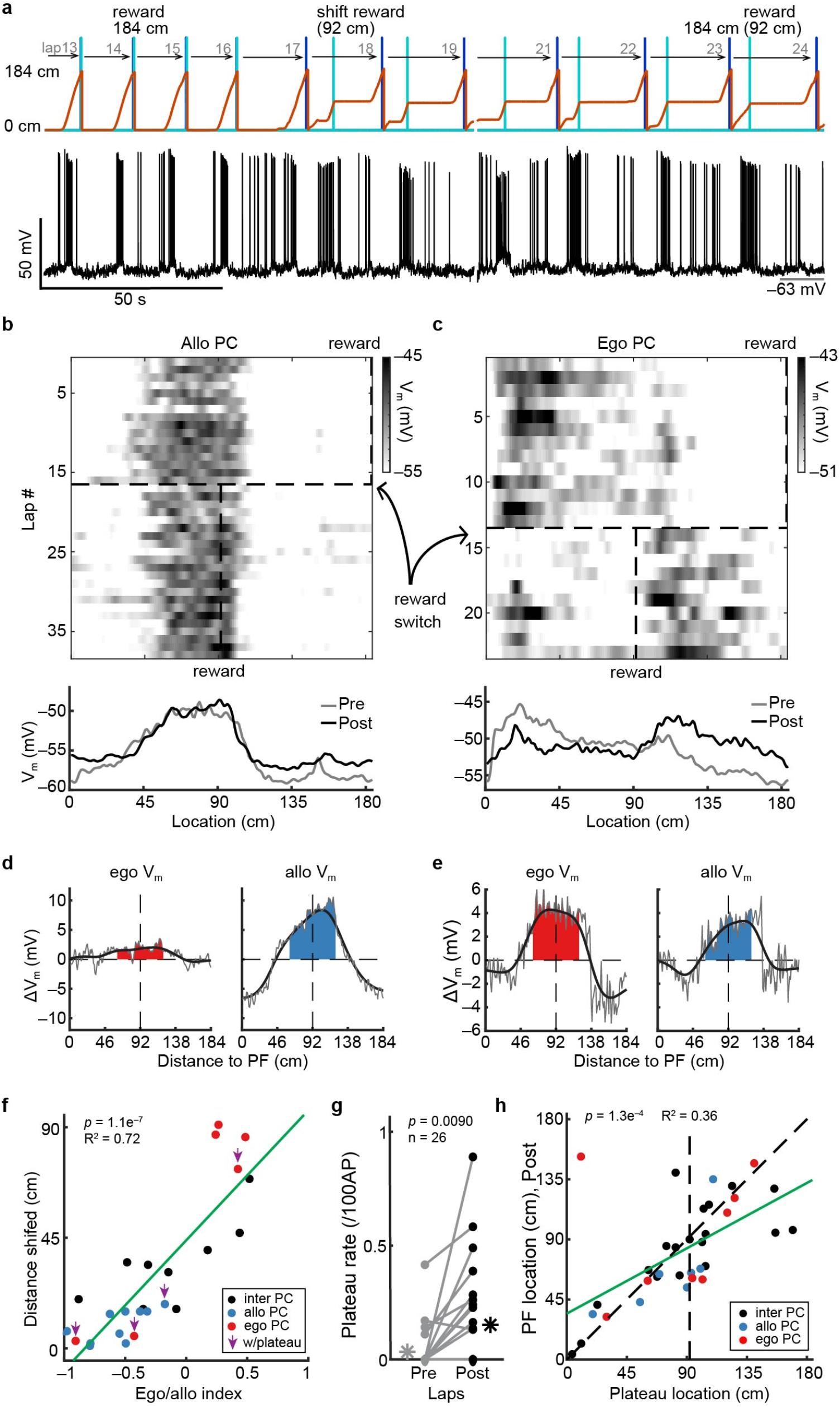
Individual PCs receive both ego- and allocentric referenced synaptic inputs. **a,** Example Vm trace of an allocentric PC in area CA1 (black), location of the mouse on the belt (red, from 0 to 184 cm), reward location (light blue line at 184 and 92 cm during Pre and Post, respectively), and the end of the lap (dark blue line, 184 cm). Pre: before reward switch; Post: after reward switch. **b,** Data from a representative allocentric PC during reward switch (same cell as in **a**). Top: shaded heatmap shows the Vm in space across laps, dashed vertical lines indicate the reward locations (at 184 and 92 cm during Pre and Post, respectively). The dashed horizontal line marks the trial of reward switch. Bottom: the averaged Vm traces in Pre (grey) and Post (black). **c**, Same as **b**, but a representative egocentric PC. **d**, ΔVm produced by reward switch for Vm traces referenced to the space (red, ego Vm) or to the start of running (blue, allo Vm), corresponding to the allocentric PC in **b**. **e**, Same as **d**, but the ego Vm and allo Vm corresponded to the egocentric PC in **c**. **f**, The correlation of ego/allo index and the distance of PF shift after reward switch but before spontaneous plateaus. The arrows indicate the cells that changed their categories due to spontaneous plateaus. The black, blue and red dots indicate the intermediate, allo- and ego-centric cells, respectively. The green line is a linear fit (n = 25 cells, *p* = 1.1 × 10^−7^, R^2^ = 0.72). **g**, The plateau rate per 100 spikes in Pre (grey) and Post (black). The far left and right asterisks are the average (n = 26 cells, *p* = 0.0090, paired-sample *t*-test). **h**, The correlation of spontaneous plateau locations and the PF positions after the given plateaus. The black, blue and red dots indicate the intermediate, allo- and ego-centric cells, respectively. The green line is a linear fit (n = 35 identified plateaus, *p* = 1.3 × 10^−4^, R^2^ = 0.36).

In general, we observed a ∼90 cm shift of the V_m_ ramps of egocentric PCs (Fig. 4c), whereas the V_m_ ramps of allocentric PCs remained at the original PF location (Fig. 4b). We did, however, also notice that in most neurons the shape of the V_m_ ramp was altered after reward switch as if the synaptic input underlying the PF was composed of separate components, one that shifted (egocentric input) and another that did not (allocentric input) (Fig. 4d–e; Extended Data Fig. 9). To quantify the ratio of these egocentric and allocentric inputs, we calculated a Λ1V_m_ under two different referencing conditions (Fig. 4d–e, Extended Data Fig. 10). In the first condition the V_m_ was referenced to the physical locations of the track (0 cm belt location; allocentric frame, Extended DataFig. 10c–e) and in the second the V_m_ was referenced to the beginning of running on each lap following reward consumption (egocentric frame, Extended Data Fig. 10g–i). In the second condition the V_m_ is essentially rotated 90 cm with respect to the first condition since the running profile is shaped by the reward location which has itself been rotated 90 cm (see Methods). In both referencing conditions, we subtracted the mean V_m_ traces calculated from all the trials before the reward switch and after any spontaneous plateaus (see Methods, ≥ 4 trials, 15.08±0.63 trials) from the mean V_m_ calculated for all the trials after the switch and before any spontaneous plateaus (see Methods, ≥ 3 trials, 19.88±2.48 trials) for all PCs (ΔV_m_, n = 25 cells). The egocentric component of the V_m_ was determined as the amount of depolarization (area) that shifted 90±30 cm away from the original PF location for the traces referenced to location (allocentric frame, Extended Data Fig. 10e–f; Extended Data 10a, ego V_m_). The allocentric component was determined as the amount of depolarization (area) that shifted 90±30 cm for the traces referenced to the running (egocentric frame, Extended Data Fig. 10i–j; Extended Data 10a, allo V_m_). Summation of the two resulting components adequately reconstructed the original V_m_ in most cells, suggesting that this was a fairly accurate measure (mean deviation: 0.29±0.037; Extended Data Fig. 12).

We calculated an Ego/Allo index ((E–A)/(E+A)) from these quantities and found a strong correlation between the amount the PF shifted and the index, with egocentric PFs showing positive values and allocentric PFs showing negative values and intermediate PF with values in between these extremes (Fig. 4f; Extended Data Fig. 11b). The location-dependence of the ego/allo index was consistent with that observed in the imaging data (Extended Data Fig. 8e). Together, these results suggest that CA1 PCs simultaneously receive both egocentric and allocentric referenced synaptic inputs and that the relative proportion of ego/allo inputs is a determinant of the distance PFs shift. In addition, a level of imperfect stability or shifting also may contribute to the PF locations of some intermediate PCs (Extended Data Fig. 11c). Thus, individual PCs in CA1 receive a variable proportion of both egocentric and allocentric synaptic inputs, which largely dictate the initial remapping following a reward switch.

### BTSP also determines some place field locations

In our V_m_ recordings approximately 46 % of PCs (12 of 26) exhibited spontaneous long-duration plateau potentials^61,68^ that altered the location of the PFs by rapidly overwriting the current synaptic weights (see examples in Extended Data Fig. 13)^69^. Indeed, the plateau potential rate significantly increased after the reward switch (from 0.003 to 0.15 per 100 spikes; *p* = 0.0090, paired *t*-test; Fig. 4g), suggesting an enhanced level of BTSP could also play a role in PC remapping (Fig. 4h). To further explore how BTSP modulates PC remapping, we identified putative BTSP events from the above population imaging data recorded under familiar conditions^37,40,70^ (Fig. 5a–d; see Methods). With our criteria, we found BTSP events after the reward location switch in 63.5 % of allocentric PCs and 74.1 % of egocentric PCs. In most allocentric PCs with BTSP events (69.7 %), the events were clustered around the original PF location where the apparent plasticity induction enhanced the allocentric selectivity (Fig. 5e, Left; 5f, blue; 5g, Left; 5h, blue). In the remaining allocentric PCs with BTSP events (30.3 %), the plasticity was primarily responsible for producing the allocentric referencing as the activity of these neurons had become nonspecific or egocentric initially after reward shift (Fig. 5g, Left). For egocentric PCs with BTSP events, the events were preferentially clustered nearly 90 cm away from the original PF location (Fig. 5e, right; 5f, red). In approximately three-fifths of egocentric PCs with BTSP events (60.9 %), BTSP in a single trial shifted the PF location to produce egocentric activity while it enhanced the level of egocentric selectivity already present in the remaining 39.1 % of egocentric PCs (Fig. 5f, red; 5g, right; 5h, red). Together, the above data suggest that ego- and allo-centric referenced synaptic inputs support a conjunctive map in CA1, with appropriate shifting of these synaptic inputs and subsequent BTSP induction both contributing to the remapping observed following reward location shift.

**Fig.5.**
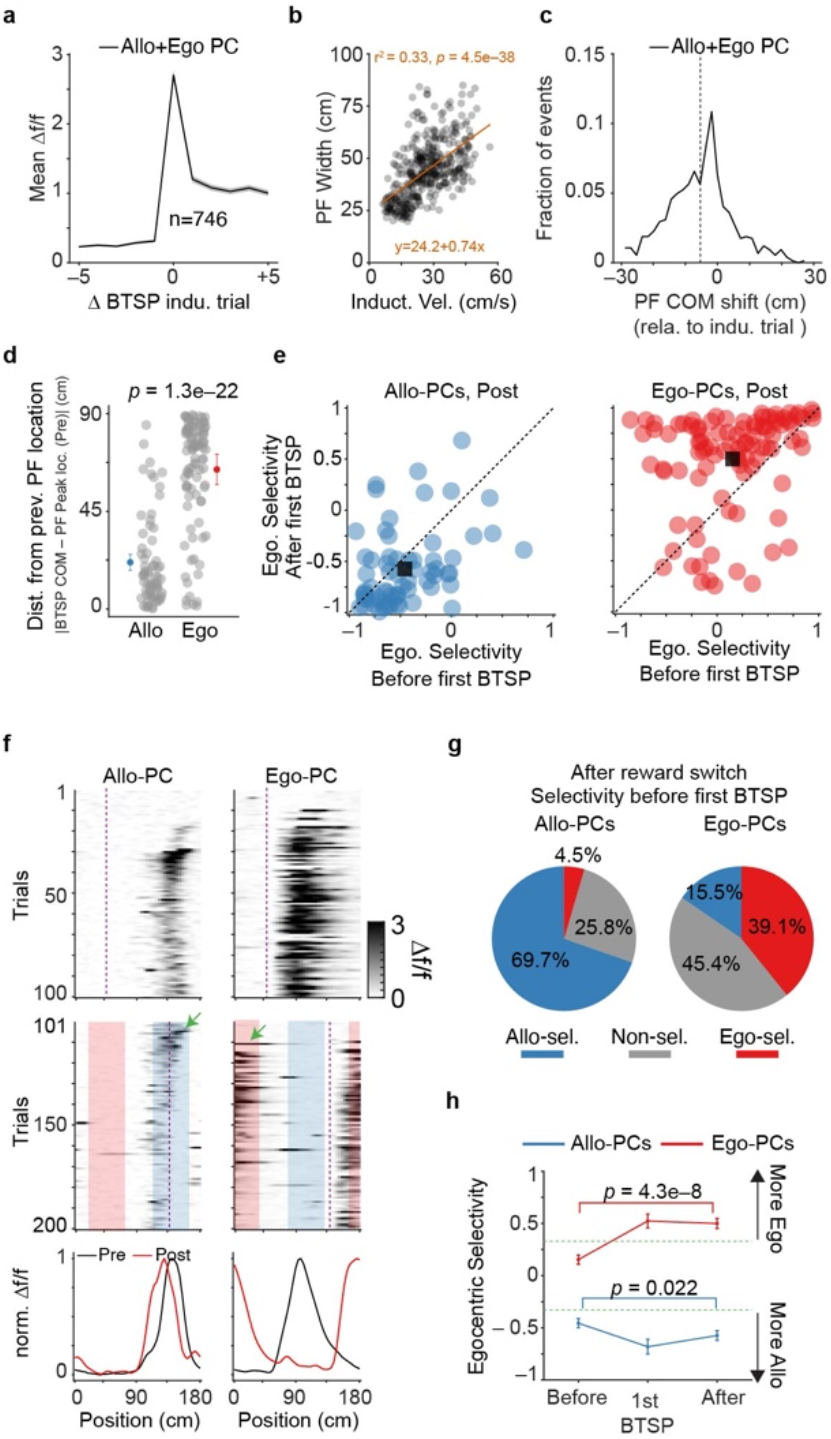
BTSP was involved in PC remapping. **a,** Mean Δf/f within field as a function of trials aligned to the BTSP induction trial before (Pre) reward switch (n = 746 BTSP events from 6 mice; see methods). Solid line and shaded areas indicate mean±SEM. **b,** Resulting PF width following BTSP as a function of running velocity during the BTSP induction event. Each dot represents one BTSP event. Orange line is a linear fit. **c,** Distribution of events as a function of the shift distance between the resulting PF COM and the COM of BTSP event. Black dashed vertical line marks the median PF COM shift. Data pooled from ego- and allo-centric PCs (235 PCs) for **a**–**c**. **d,** Distance of the first BTSP event in Post from the cell’s PF location in Pre (n = 6 mice) for allo- (left) and egocentric (right) PCs. Each dot represents one BTSP event. (Allo-PC: 21.5±2.6 cm; Ego-PC: 64.3±2.5 cm; *p* = 1.3e–22, unpaired *t*-test). **e,** Scatter plots of egocentric selectivity before and after the first BTSP events during Post for allo- (left, blue) and ego-centric (right, red) PCs. The black square marks the averaged mean egocentric selectivity. Dashed line represents unity. Those events that increased the egocentric selectivity will be located above the unity line, and similarly, those events that decreased egocentric selectivity will be located below unity. Egocentric selectivity is defined as the difference of mean Δf/f within the egocentric field from the allocentric field divided by their sum, which has a range of [–1, 1]. **f,** Example of one allocentric (left) and one egocentric (right) PC. Shaded maps depict Δf/f in space during Pre (top) and Post (middle). Green arrows mark the first BTSP events after reward switch. Dashed lines indicate the reward locations. Blue shaded area indicates allocentric field (∼45 cm wide around the original PF location in Pre), red shaded area indicates egocentric field (∼45cm wide that is 90 cm away from the original PF location in Pre). Post: after reward switch. Bottom: Peak-normalized (norm.) mean Δf/f during Pre and Post for each cell. **g,** Fractions of each type of activity prior to the first BTSP event in Post for allo- (left) and ego-centric (right) PCs (n = 6 mice). Criterion for different types: > 0.33 is ego-selective, < –0.33 is allo-selective, in between is non-selective. **h,** Egocentric selectivity before, during and after the first BTSP events from all allo- (blue) and ego-centric (red) PCs in Post. (Allo-PC: before is –0.46±0.044, after is –0.57±0.046, *p* = 0.022, Paired *t*-test; Ego-PC: before is 0.15±0.045, after is 0.50±0.049, *p* = 4.3e–8, Paired *t*-test). Green dashed lines mark two thresholds. All data are from the familiar belt, first day of reward switch (n = 6 mice, same animals as Fig. 1).

## Discussion

### Summary

Here we report that PCs with both egocentric and allocentric reference frames contribute to a conjunctive spatial map of a familiar environment. We find this conjunction at both the population level (egocentric and allocentric PCs) and the cellular level (egocentric and allocentric synaptic inputs to PCs). We also report that exposure to a novel environment reshapes the conjunctive map with individual PCs flexibly changing reference frames mainly from allocentric to egocentric. This dynamic PC remapping in CA1 appears to be mediated by a combination of factors such as alterations in the location-dependence of synaptic input, perhaps explained by changes in the activity of area CA3, and new synaptic plasticity in CA1 rapidly produced by dendritic plateau potentials (BTSP).

### A conjunctive spatial map in CA1

We found that when an animal is engaged in a goal-directed spatial learning task in a familiar environment, a conjunctive cognitive map in the hippocampal CA1 is instantiated by a mixture of egocentric and allocentric referenced PCs. Moreover, egocentric PCs tend to cluster after the reward site, suggesting that the CA1 representation is primarily egocentric at the beginning of each lap, consistent with previous work^32–34^. Notably, the majority of egocentric PCs have their PFs more than 30 cm in space and 6 seconds in time following the reward site, suggesting that it is unlikely that these egocentric PCs represent direct reward responses. The past reward site may however function as an egocentric starting position to update an animal’s present position primarily through the process of path-integration^5,10–12^. As the animal moves further away from its starting position, we show that the PC representation relies more heavily on allocentric spatial cues which can, in principle, correct the path integration drift error^10–12^. Notably, the fraction of allocentric PCs has a strong inverse correlation with that of egocentric PCs, suggesting a competitive relationship^32^.

### Experience-dependent flexible referencing

After mice have learned the spatial relationship between the reward and the spatial cues in a familiar environment, performing the same task in a novel environment covered with unfamiliar external cues causes the PC representation to become predominately egocentric. Moreover, our longitudinal recordings confirm that a large fraction of allocentric PCs adaptively become egocentric PCs in this process. A parsimonious interpretation is that, under our experimental conditions, the animal may largely disregard the unfamiliar cues on the novel belt and registers its current location solely by the distance from its last starting position through path integration. This is consistent with previous work suggesting that egocentric reference frames alone are sufficient to maintain an established map^8,17,24,25^. Overall, our results clearly demonstrate that the anchoring of individual PCs to either allocentric or egocentric reference frames is not fixed but rather flexibly shaped by experience.

### Synaptic mechanisms

Previous studies revealed that dendritic plateau potential driven BTSP adjusts the synaptic weights of inputs to produce the large V_m_ ramps that drive PF firing in CA1^6,61–63,69^. Our whole-cell recordings suggest that the majority of synaptic inputs with strong weights on CA1 PCs remain active following a reward location change with these inputs either maintaining or immediately altering their location-dependence. This suggests that new synaptic plasticity through BTSP is not necessarily required for the initial remapping process in many PCs. Rather, our data suggest that much of the immediate PF reorganization in CA1 after reward switch is inherited from its upstream regions, presumably CA3^60,71,72^. Interestingly, V_m_ ramps of most PCs split into egocentric and allocentric components, indicating that individual CA1 PCs simultaneously receive both egocentric and allocentric referenced synaptic inputs to produce their conjunctive PFs. The proportion of potentiated egocentric and allocentric synaptic inputs that each PC receives then primarily determines the degree of PF shifting in these PCs.

We also find, however, that the reward switch elevates the rate of plateau potential initiation in CA1 and that this rapidly reshapes the PF referencing of many PCs by overwriting the synaptic weight matrix through BTSP. Further, our data suggest that BTSP might preferentially enhance the PFs of some allocentric PCs and may also abruptly form a large fraction of the ego-centric PC population. Together, we propose that the reorganization of allocentric and egocentric referenced feedforward CA3 synaptic inputs and the induction of new BTSP together produces the PC remapping observed in CA1 after reward location switching.

## Conclusion

Understanding how different reference frames are encoded and retrieved in the hippocampus could ultimately help improve our understanding of how a cognitive map is formed and retrieved there^1,2,5^. It is commonly believed that the integration of egocentric and allocentric reference frames is hierarchical across brain regions, and the representation follows an egocentric-to-allocentric trend from the parietal cortex to the retrosplenial cortex and then to the hippocampus^3,22,23,73,74^. Our observation that CA1 PCs are formed by simultaneous input from both egocentric and allocentric referenced synaptic activity suggests instead that these reference frames are integrated, perhaps for the first time, within CA1. But this remains an open question, as future experiments are needed to determine PC referencing in upstream regions particularly in area CA3. Finally, that the degree of this reference frame mixing is dependent upon the experience of the animal suggests that mechanisms exist to link the behavior of the animal with the individual components of the cognitive map^55,75^.

## Supporting information

FlexibleEgoAlloCA1_supFig

## Acknowledgements

We thank R Chitwood for technical support, the Magee lab members for useful comments on the manuscript. This work was funded by HHMI (JCM), the Cullen Foundation (JCM) and HHMI via Life Sciences Research Foundation (YL).

## Author contributions

Conceptualization: FKQ, YL, JCM

Two-photon imaging recording and data analysis: FKQ

Whole-cell recording: YL

Whole-cell data analysis: YL, JCM

Writing, review & editing: FKQ, YL, JCM

## Competing interests

No competing interests to be declared.

## Data and materials availability

The data supporting this study’s findings are available from corresponding author upon request. The code that supports, the results of this study will be made available via a GitHub repository.

## Methods

All experimental procedures performed were approved by the Baylor College of Medicine Institutional Animal Care and Use Committee.

### Surgery

All experiments were performed in adult GP5.17^39^ (n = 22 mice, at least 8 weeks at the day of surgery) of either sex or WT male C57BL6 (n = 16 mice, 8–12 weeks old). The experimenter was not blind to different conditions of the experiments. Animals were housed in the Magee Satellite under an inverse 12-hour dark/12-hour light cycle (light time: 9 pm–9 am), with controlled temperature (around 21°C) and humidity (around 30–60 %). All recordings were carried out starting at the dark cycle. All surgeries were performed with mice anesthetized under ∼2 % isoflurane. The surgery procedure was performed as described previously^37^. After the application of local anesthetics, a small flap of scalp skin was removed to expose the skull, the skull was then cleaned. We then levelled the skull and marked the craniotomy center for CA1 imaging targeted at (in mm) 2.3 (AP, posterior to bregma) and 2.15 (ML, lateral to midline); while at 1.9 (AP) and 2.0 (ML) for CA1 intracellular recordings (right hemisphere). The surgery for intracellular recordings was identical to previous detailed reports^6, 61^. For two-photon imaging, a 3.0-mm-diameter craniotomy centered at the coordinate described above was then created, dura was removed using forceps (Fine Science Tools), then the overlying cortical tissue was gently aspirated using a blunt needle (McMaster-Carr) under repeated irrigation. Aspiration continued to expose the external capsule, then the external capsule was lightly peeled away without the suction needle directly contacting the fibers. Saline irrigation continued until the bleeding around the center of the field of view stopped. A cannula (3 mm diameter, 1.75–2.0 mm long) with cover glass (Potomac, 2.90 mm diameter, #0) glued on the bottom using UV-curable optical adhesive (Norland Products) was inserted and cemented. Cannulae were tightly fit with the craniotomy for better stability. Dental cement (Ortho-Jet, Lang Dental) was then applied to attach a custom-made titanium head bar to the skull parallel to the surface of the cover glass to acquire optimal recording quality.

### Behavior task (two-photon imaging)

The linear treadmill system consists of a velvet fabric made belt (McMaster Carr) that is ∼180 cm long with mice head-fixed under the head posts. The belt was self-propelled by water restricted mice, the movement speed of the belt (therefore the animal) was measured by a rotary encoder using Arduino-based microcontrollers, the digitized movement speed signal interfaced with a behavior control system through a Bpod module (Bpod r0.9–1.0, Sanworks) using MATLAB custom code (2019b, MathWorks) running on a Windows PC. The reward delivery of 10 % sucrose water is controlled by a solenoid valve (The Lee Co.) through a custom-made lick port. The animal’s licking was detected by a custom fabricated optical lickometer. The Bpod system interfaced with the rotary encoder, the valve, and the sensor. Behavior data were digitized through a PCIe-6343, X series DAQ system (National Instruments) and saved via WaveSurfer software (v0.982, Janelia).

Animals are allowed at least 6 days to recover from the optical window implantation surgery before being placed on water restriction (1.5 ml/day). Animals were then handled by the experimenter for at least 10–30 min, for at least 4 days (usually 5–6 days). We then introduced the treadmill system to the head-fixed mice for 3–5 days. Mice were then trained on the featured belt to run for a 10 % sucrose water delivered following epoch-to-epoch increasing distance. Each epoch has 8–15 laps. The distance between rewards is fixed within each epoch and increased by epoch. Manual rewards were released by the experimenter to encourage running during the early phase of training. Once the distance between rewards reached 126 cm, the reward was then delivered at a fixed location (50 cm) on the belt. Animals were allowed to run for at least 4 days (typically 6 days, 700 laps in total) with fixed reward location before we started the reward switch experiment. The last day of fixed reward experiment was named ‘day 0’. We then started to perform the reward switch experiment, where the reward was fixed at 50 cm for the first half of session (100 trials), then the reward was switched to 140 cm for the second half of recording session (100 trials). The first day of reward switch was named ‘day 1’. Individual sessions typically last for 30–60 min, with one recording session per day. Two belts are used: Belt A uniformly covered by 3 cues (Velcro patched, glue sticker, white fabrics, each cue region is 60 cm). Belt B uniformly covered by 6 cues (different from Belt A, each cue region is 30 cm).

### Two-photon imaging

All two-photon Ca^2+^ imaging recordings were performed in the dark using custom-made microscope (Janelia MIMMS 2.0 system). Transgenically expressed GCamp6f in the hippocampal CA1 was excited at 920 nm (typically 30–50 mW, measured through the objective) by a Ti:Sapphire laser (Cameleon Ultra II, Coherent) and imaged through a Nikon 16×, 0.8 Numerical Aperture (NA) objective. The emission light passed through a 565 DCXR dichroic filter (Chroma) and a 532/46 nm bandpass filter (Semrock) and was detected by a GaAsP photomultiplier tube (11706P-40SEL, Hamamatsu). Images (512 × 512 pixels) were acquired at ∼30 Hz using the ScanImaging software (Vidrio Technologies, LLC).

Imaging field of views (FOVs) (∼300 × 300 µm) were chosen by visual examination of the presence of Ca^2+^ transients in the somata. For longitudinal imaging, a reference FOV was first chosen and registered. Each day, the FOV was aligned to this reference. Experiments will be aborted if significant differences were noticed on subsequent consecutive days in the imaging FOV.

### In vivo intracellular recordings

The intracellular recordings were performed as previously reported^6, 61^. Briefly, an extracellular LFP electrode was firstly lowered into the dorsal hippocampus using a micromanipulator until prominent theta-modulated spiking and increased ripple amplitude were detected after passing through neocortex, usually to a depth of 1.0–1.2 mm. Then a glass intracellular recording pipette was lowered to the same depth while applying positive pressure (∼9.5 psi). The intracellular solution contained (in mM): 134 K-Gluconate, 6 KCl, 10 HEPES, 4 NaCl, 0.3 MgGTP, 4 MgATP, 14 Tris-phosphocreatine, and 0.2 % biocytin. Current-clamp recordings of intracellular membrane potential (V_m_) were amplified and digitized at 20 kHz, without correction for liquid junction potential.

## Data analysis

### Pre-processing of imaging data and signal processing

Acquired two-photon images were motion-corrected using Suite2p^76^ (Python version, http://github.com/MouseLand/suite2p). For longitudinal imaging across multiple days, all acquired data from each day were concatenated and motion corrected. Regions of interests (ROIs) were selected, time series fluorescence traces were extracted automatically using Suite2p. Manual curation of ROI selection was performed to discard unwanted ROIs and re-select ROIs that were automatically discarded by Suite2p based on the anatomy in the Suite2p Graphical User Interface (GUI). Next, ROI-to-ROI signal contamination was examined using the Suite2p GUI as followings: ROIs that are physically close were selected (typically 10–20), and the time-series fluorescence traces were examined visually to identify all possible ‘ROI-x contaminates ROI-y’ combinations. Two rounds of examinations were performed for each FOV, and all possible contaminations were examined. Contaminated ROIs were marked and then discarded at the end of each round. Typically, ∼40 % ROIs were discarded after 2 rounds of examinations. No neuropil subtraction was performed. Further analyses were then performed using custom codes written in MATLA (version 2021a). Briefly, the raw fluorescence was first baseline-corrected using a 5000-frame window before converted to Δf/f, which was calculated as (F–F_0_)/(F_0_), where F_0_ was the mode of the histogram of F. Significant calcium transients were defined as transients larger than 3 times standard deviations above the baseline. Standard deviations of the baseline noise were estimated from deviations below peak histogram values of all Δf/f activity. Spatial map of Δf/f was then generated for each ROI using only the frames where animals’ velocity is larger than 5 cm/s. The belt length (∼180 cm) is divided uniformly into 50 spatial bins (3.6 cm/bin), and the mean Δf/f was calculated for each spatial bin. A gaussian smoothing of 3 spatial bins was applied to each spatial map of Δf/f for display purpose. To have a consistent lap number across animals and conditions, only the first 100 trials before, and the first 100 trials after, reward switch were used within the same conditions. The reward is either at bin 14 (50 cm, before switch) or bin 39 (140 cm, after switch). All day 0 recording sessions have at least 100 laps. If one condition has less than 100 laps, we analyzed all the available laps. The reward position was marked for every presented figure.

### Place cell identification

We first determined the onset trial of each PC as the first trial where: 1) at least one spatial bin had significant events (larger than 3* SD above the baseline); 2) at least 2 out of 5 of the following laps had at least one spatial bin with significant events. The significant event was detected within 90 cm (±12 spatial bins) around the peak position of the trial-averaged place field activity. Moreover, the max, but not the mean, Δf/f within each spatial bin was used to determine if any spatial bin had significant events detected. Only the first onset lap was selected for each cell during one condition (e.g., reward at one location). Only the trials subsequent to the onset lap were used to determine if a cell is a PC.

A cell is defined to have spatially modulated activity if the trial-averaged PC activity provided enough spatial information about the linear track locations (larger than 95^th^ percentile of the shuffled spatial information). The spatial information (SI) for each cell is calculated as previously described^31^:

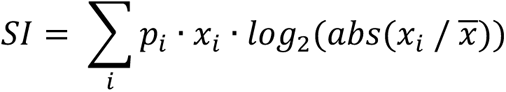

Where *p_i_* is the probability of occupancy for spatial bin *i*, *x_i_* is the smoothed mean activity (Δf/f) while occupying bin *i*, *x̅* is the overall mean activity level. The SI for each cell is then compared to 200 shuffles of the activity (each shuffle was generated by circularly shifting the Δf/f time series traces by at least 500 frames, then dividing the trace into six trunks and permuting their order). If the observed spatial information exceeded the 95^th^ percentile of the shuffled information values, its field was considered spatially modulated. A cell is defined as reliable if more than 20 % of the trials (after the onset lap) had significant events. A cell is defined as a PC if the cell is both spatially modulated and reliable. If a cell doesn’t have any identified onset lap, onset is set as ‘NaN’ but the identification of PC for this cell continued using all trials starting from the first trial. If one cell had multiple place fields, only the stronger place field (defined as the peak from the average place field activity) was considered and no further processing was added.

### Spatial map of the behavioral data

Same as the spatial map of Δf/f analysis, the spatial maps of running velocity and counts of licks for each animal were generated by dividing the belt length (180 cm) into 50 spatial bins with each spatial bin as 3.6 cm. Then, the mean velocity (cm/s) and the total lick count were generated within each spatial bin. Data when velocity is below 5 cm/s were discarded before the analysis. The lick probability trace for each animal was calculated as the fraction of trials that had at least one lick within each spatial bin.

### Identification of start of running

Briefly, we identified the running onset as follows. The velocity and distance data are sampled in 10k Hz. From the time of reward delivery, we first identified the First low-speed point within the next 5 seconds that has speed < 1cm/sec. If no such time point was found, we then changed the criterion to 5 cm/sec. If again no such time point was found, we took the lowest speed point within 0.5 seconds after reward delivery as the First low-speed point. The distance was then realigned so that the realigned distance around the First low-speed point was zero. We next calculated, using a 1-second sliding time window, the fraction of time that animal had speed smaller than 1 cm/sec. The first time point where this number is 0.5 (meaning 0.5 seconds out of 1 second animals had < 1 cm/sec speed) was taken as the onset of running. If no such onset of running was identified, we changed the low speed threshold to 5 cm/sec, and searched again. If again no such onset of running was identified, we searched within the First Low Speed Point and the time where realigned distance reached 18 cm for the last time where speed is < 5 cm/sec. If such last low speed point was found, we identified the first time after the First low-speed point where realigned distance is > 5 cm as the onset of running. If such last low speed point was not found, we picked the lowest speed point between low-speed Point and the time where re-aligned distance reached 18 cm. In extreme cases where none of the above scenarios were met, the reward delivery time was taken as onset of running. Mice typically stopped for more than 5 seconds, and our search constraints were merely designed for extreme cases. Every trial in Pre was included to calculate the averaged time between reward delivery and the onset of running. Only the first 100 laps before reward switch on day 1 were analyzed (n = 6 mice).

### Determination of place cell types

Under our experimental conditions, we defined ‘Allo’ (allocentric) PCs as those that maintain stable PFs relative to fixed spatial cues on the belt ((∼15 cm shift), and ‘Ego’ (egocentric) PCs that shifted PFs following the new location of the reward (75–90 cm), and ‘Inter’ (intermediate) PCs that had shifted 15–75 cm. See Extended Data Fig. 2a for the identification of criterion.

### Population vector correlation analysis

To understand how the PC population encodes each location, we used population vector analysis ^54^. Briefly, population vectors (PVs) refer to activity vectors at each location where each element from the vector represents the activity level of a specific neuron. The Pearson correlation between the population vectors across all locations gives rise to the similarity matrix *M*. Each element *M*_!,$_ represents the Pearson correlation coefficient between population vectors at location *i* and location *j* on the linear track. Each linear track is divided into 50 spatial bins, therefore the size of *M* is 50 by 50. The similarity matrix therefore provides a measure of how similar the representation of a given position is between different conditions. When comparing PVs before versus after the reward switch, a few possibilities to be emphasized: 1) if the PC representation is completely allocentric, and the reward switch will have no effects on the PC representation, then a high-similarity region along the diagonal is expected. 2) if the PC representation is completely egocentric, and the PC representation exclusively encodes position information related to the run start, then a high-similarity region along the off-diagonal (a 90 cm shift) is expected. 3) If the relative dominance by allocentric and egocentric PCs had a clear location preference (e.g., egocentric PCs predominantly clustered after the reward location, the rest of track was dominated by allocentric PCs), then coherent high-similarity region along neither the entire diagonal nor the entire off-diagonal band would be observed. Instead, fragmented smaller high-similarity patches could be observed as some aligned to the diagonal (more allocentric) and others aligned to the off-diagonal (more egocentric). 4) If Intermediate, but neither allocentric nor egocentric, PCs dominate the PC representation, and the Intermediate PCs at the same location tended to shift similar distances, then a high-similarity region along the areas between the diagonal and off-diagonal was expected. To quantitatively compare the influences of allocentric spatial coding and egocentric spatial coding, we averaged along the diagonal elements of the similarity matrix *M* to represent ‘allocentric similarity’, and we averaged along the off-diagonal elements of the similarity matrix *M* as the ‘egocentric similarity’.

### BTSP events identification

Our analysis on BTSP event identification was similar to previous studies with some modifications^37, 40, 70^. We identified putative BTSP events as 1) a ‘strong’ significant Ca^2+^ event with its amplitude in the top 10^th^ percentile for all significant Ca^2+^ event amplitudes identified within the same session in the same cell. Laps before and after the reward switch are used to measure their separate thresholds for ‘strong’ activity. 2) Only trials with peak Δf/f within 12 spatial bins (43.2 cm) of putative BTSP event peak were considered for BTSP identifications. 3) four out of five of the subsequent trials had significant Ca^2+^ event. 4) The amplitude of the following 5 trials increased at least 100 % compared to that of the preceding 5 trials, and 5) the identified BTSP events had peak Δf/f larger than 2. 6) We enforce that the last five laps (lap 96–100, lap 196– 200) do not have BTSP events due to lack of sampling laps, and releasing this criterion doesn’t change our conclusions. Moreover, the max, but not the mean, Δf/f within each spatial bin was used to determine if any spatial bin had significant events detected. Significant events were identified if the amplitude was above our previously identified significant event threshold (Mean + 3*SD). For identified significant Ca^2+^ events, PF of each event was quantified as the consecutive spatial bins where max Δf/f exceeded 35 % of the peak of max Δf/f value. The total number of such spatial bins was then the PF width of this event. The PF of the mean of the subsequent or preceding 5 trial activity was defined as the following^77^: 1) consecutive spatial bins where max Δf/f exceeded 35 % of the peak of max Δf/f values. 2) PF width smaller than 90% of the entire track. 3) In-field max Δf/f larger than 0.2. 4) The mean in-field Δf/f must be greater than 2 times of mean of out-field Δf/f. The Center of Mass (COM) of the identified BTSP events was calculated as the following equation:

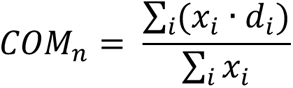

Where *x_i_* is the max Δf/f within spatial bin *i*, *d_i_* is the spatial position from the start of the track. COM for any given event only used the within PF spatial bin data as described above. The COM of the mean of the subsequent or preceding 5 trials was calculated the same way as described above if a PF was found. Induction velocity of any events was calculated as the mean of the velocity within the identified PF.

### Intracellular recordings

To analyze the spatial location of AP rate, the spatially binned AP rate was determined (AP#/time in bin) using the spatial bin size of 1 cm for each trial and averaged over the duration of the recording. To analyze V_m_ ramps, APs were removed, baseline corrected at trial basis by subtracting the difference in the recorded AP threshold (the most negative 5^th^ percentile) and –50 mV and spatially binned and averaged as above. The spatially-binned AP rates and V_m_ ramps in the heatmap were smoothed with a Gaussian of 5 bins (5 cm) and 11 bins (11 cm), respectively. The PF location was defined as the COM of AP rate (smoothed with Gaussian, and those spatial bins with firing rate < 10 % of peak firing rate were not counted as part of the mass) averaged from all the trials before and after reward switch (or before/after plateaus as stated in the figures or texts) in a given PC. The egocentric PF was defined as 92 cm away from the original (allo) PF before reward switch.

### Ego V_m_, allo V_m_ and ego/allo index

To calculate the putative egocentric V_m_ ramps, we first referenced all the V_m_ ramps to physical locations of the track (0 cm location on belt). Next, V_m_ ramps from all the trials before (after plateaus if any, ≥4 trials, 15.08 ± 0.63 trials) and after (before plateaus if any, ≥3 trials, 19.88 ± 2.48 trials) reward switch were averaged. Then the before and after reward switch V_m_ ramps were subtracted (ΔV_m_ = V_m__post – V_m__pre). The AUC (area under the curve of ΔV_m_ ramps) that covered ±30 cm around the egocentric PF location (red window in Extended Data Fig. 10e, 92 cm away from original PF location), was used to define the ego V_m_. The putative allocentric V_m_ ramps were similarly calculated but with referencing of all the V_m_ ramps to the start of running before the subtraction. Thus, the allocentric PF shifted 92 cm away from the original PF location when aligned to the start of running. We defined the allocentric area as ±30 cm around the allocentric PF location, and the allo V_m_ was defined as the AUC of allocentric area. After obtaining the ego V_m_ and allo V_m_ for each individual cell, we calculated the ego/allo index as:

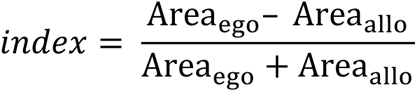

One cell was excluded from this analysis because of immediate spontaneous plateau potentials after reward switch. And the symmetry of ΔV_m_ for a given cell is:

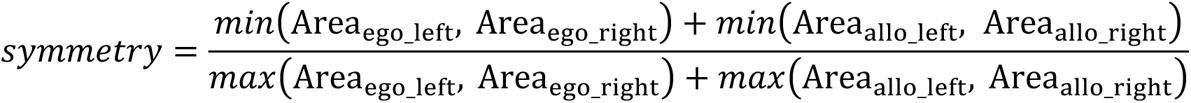

The left and right areas are the areas from –30 to 0 cm and from 0 to 30 cm relative to ego- or allocentric PF, respectively. And another 9 cells were excluded because of either the spontaneous plateau potentials (4 cells, marked with a star ‘*’) or two separate PFs appeared post-reward switch (5 cells, marked with a ‘#’ symbol). Then we reconstructed the PF V_m_ ramps by summating the ego V_m_ and allo V_m_ for each individual cell.

The above calculation of ego V_m_ and allo V_m_ has some potential limitations and should be considered approximate. For example, the V_m_ and V_m_ are calculated using the subtraction from a certain baseline (V_m_ before reward switch), and any depolarization already present in the baseline at the ego- or allocentric PF location would be subtracted thus underestimating either ego V_m_ or allo V_m_ depending on the referencing of this activity. Given that all PFs were induced by repetitive plateau initiation at the beginning of each recording, the amplitudes of these baseline depolarizations should generally be small thus producing a small error. Similarly, if the PC has a very broad field (>90 cm), the subtraction could produce a narrow peak with a broad negative ΔV_m_ on both sides (such as the cells shown in Extended Data Fig. 12a, the 3rd, 8th cell in the allo PC), which under-estimates both ego V_m_ and allo V_m_, with the ratio being less affected. Thus, we attempted to restrict the area of ego and allo within ±30 instead of ±45cm around the ego- and allo-centric PF to avoid those overlapping area at the edge. In general, the reconstructed PF V_m_ ramps fit the original PF V_m_ ramps quite well in the middle of the PF attesting to the overall accuracy of the method.

### Plateau potentials

Spontaneous long-lasting naturally-occurring plateau events were defined if the duration of a plateau potential (plateau potentials detection V_m_ > –35 mV) was longer than 100 ms, and the inter-plateau interval was longer than 10 ms. To measure the PF position before/after plateau, only those plateaus with at least 2 plateau-free trials before and after a given plateau were taken.

### Statistical Methods

The exact sample size (n) for each experimental group is indicated in the figure legend or in the main text. No statistical tests were used to predetermine sample sizes, but our sample sizes are similar to those reported in previous publications (population imaging^36, 43, 70, 78^ and in-vivo whole-cell^61, 64, 66^) using a similar behavior task and are guided by the number of neurons that can be imaged using two-photon microscopy or patched in awake behaving mice. In some cases, where data distribution was assumed, but not formally tested, parametric *t*-tests were applied to analyze the data. Experiments were randomized by randomly assigning litter mate mice to the experimental groups. Data analyses were not blind to the experimenter. However, analysis was performed automatically, without considerations of trial types or experimental groups. If not otherwise indicated in the figure, that data are shown as mean±SEM.

## Notes

### Competing Interest Statement

The authors have declared no competing interest.

