## Supplementary material for "Experience-dependent place-cell referencing in hippocampal area CA1": FlexibleEgoAlloCA1_supFig

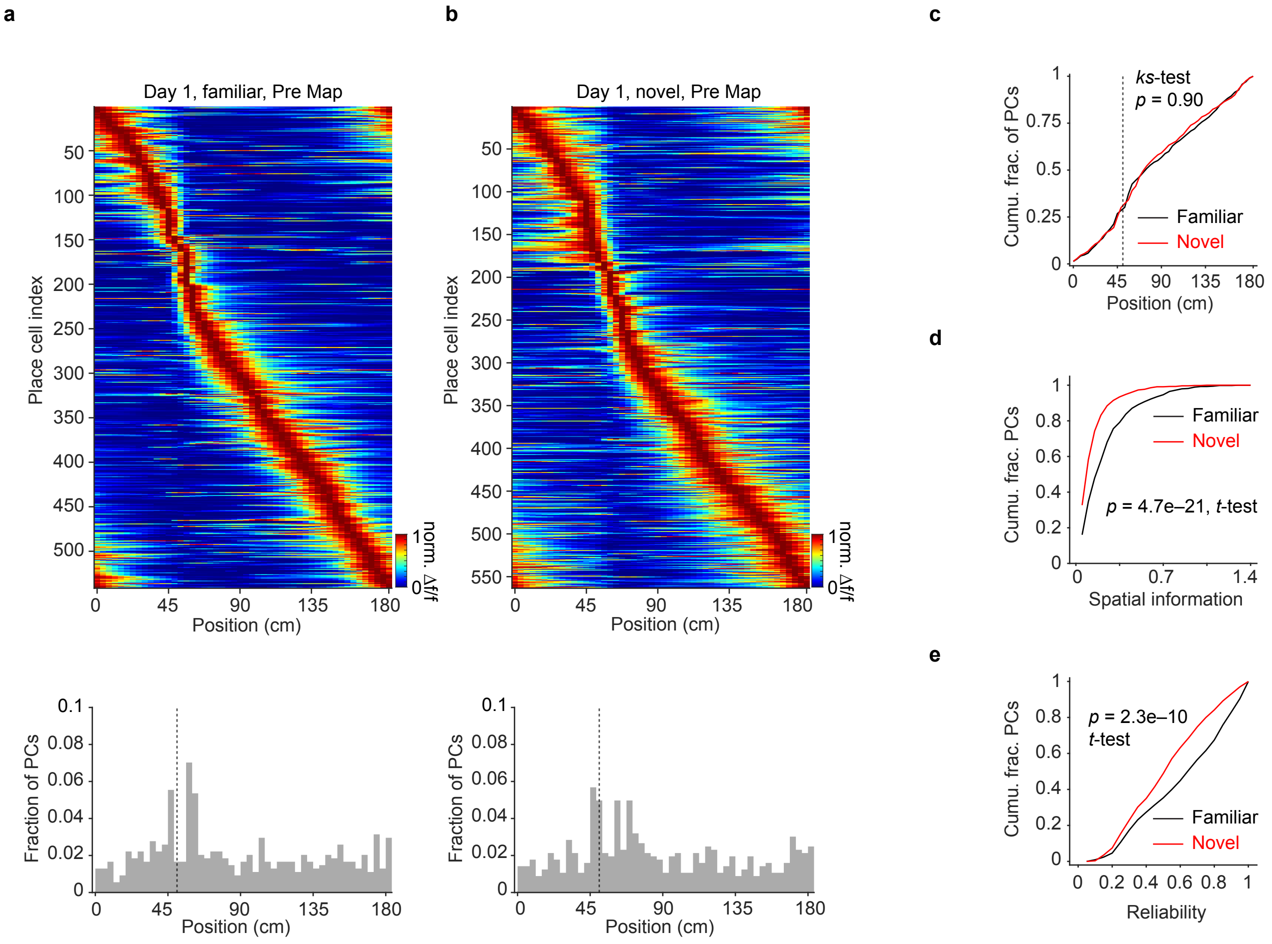


**Extended Data Fig. 1**

**Extended Data Fig. 1 | CA1 PC representation of a familiar belt versus novel belt. a,** Top: PC representation showing mean Δf/f across space including all PCs of a familiar belt, on day 1, in Pre (n = 6 mice). The PCs are ranked by Δf/f peak location. Bottom: distribution of peak position of PCs shows an over-representation near the reward location. Black dashed line marks the location of reward. Pre: before reward switch. Day 1: first day of reward switch. **b,** Same plots as **a** but for a novel belt in Pre (n = 6 mice). **c,** Cumulative fraction of PCs as a function of position for familiar and novel belt. (k-s test: Kolmogorov-Smirnov test, $p = 0.90$). **d,** Cumulative fraction of PCs as a function of spatial information on a familiar belt versus novel belt (Familiar: 0.23±0.0093; Novel: 0.12±0.0057. unpaired $t$-test, $p = 4.7e{-}21$). **e,** Cumulative fraction of PCs as a function of PC's trial-by-trial reliability (Familiar: 0.61±0.01; Novel: 0.52±0.01. unpaired $t$-test, $p = 2.3e{-}10$). Reliability is defined as fraction of trials exhibiting significant $Ca^{2+}$ event within the PF.

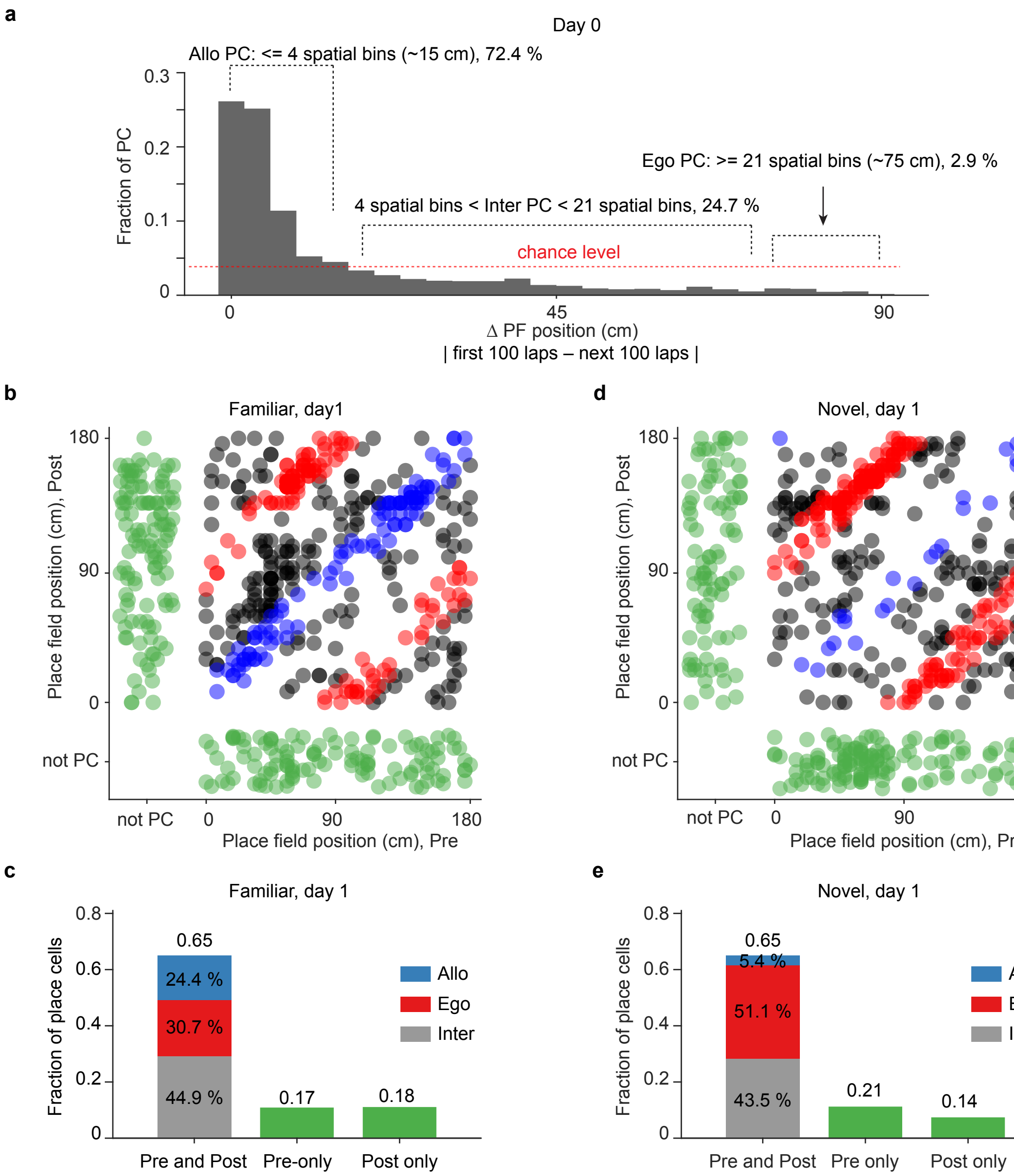


**Extended Data Fig. 2**

**Extended Data Fig. 2 | CA1 PC remapping following reward switch on a familiar versus novel belt. a,** Criterion for identifying different types of PCs. Fraction of PCs as a function of PF position change, between the first and second 100 laps, on the last day of fixed reward training on a familiar belt (day 0). n = 14 mice, pooled from all animals trained on belt A. Dashed red line marks the mean chance level. The belt length (180 cm) is divided into 50 spatial bins (3.6 cm/bin). The criterion is selected as the last bin that is above chance, which is the bin #5 (first bin is 0 cm difference). The criterion for allocentric PCs is therefore ±4 spatial bins (~±15 cm), for egocentric PCs is 90±15 cm, the remaining is defined as intermediate PCs (15–75 cm). The fractions of each type of PCs are indicated by black dash lines for validating the criterion in a constant environment with only experiential and temporal evolution (number of running laps, and approximate amount of time). **b,** Scatter plot depicts the PF position during Post versus Pre on a familiar belt on day 1 (n = 6 mice). Pre: before reward switch. Post: after reward switch. Day 1: first day of reward switch. Allo-, ego- and inter-PCs are labelled in blue, red and black, respectively. Cells only having a PF either during Pre (disappeared during Post) or Post (newly appeared during Post), but not both, are labelled in green. **c,** The fraction PCs that fall into different categories. **d-e,** Same plots as **b-c** but for a novel belt on day 1 (n = 6 mice).

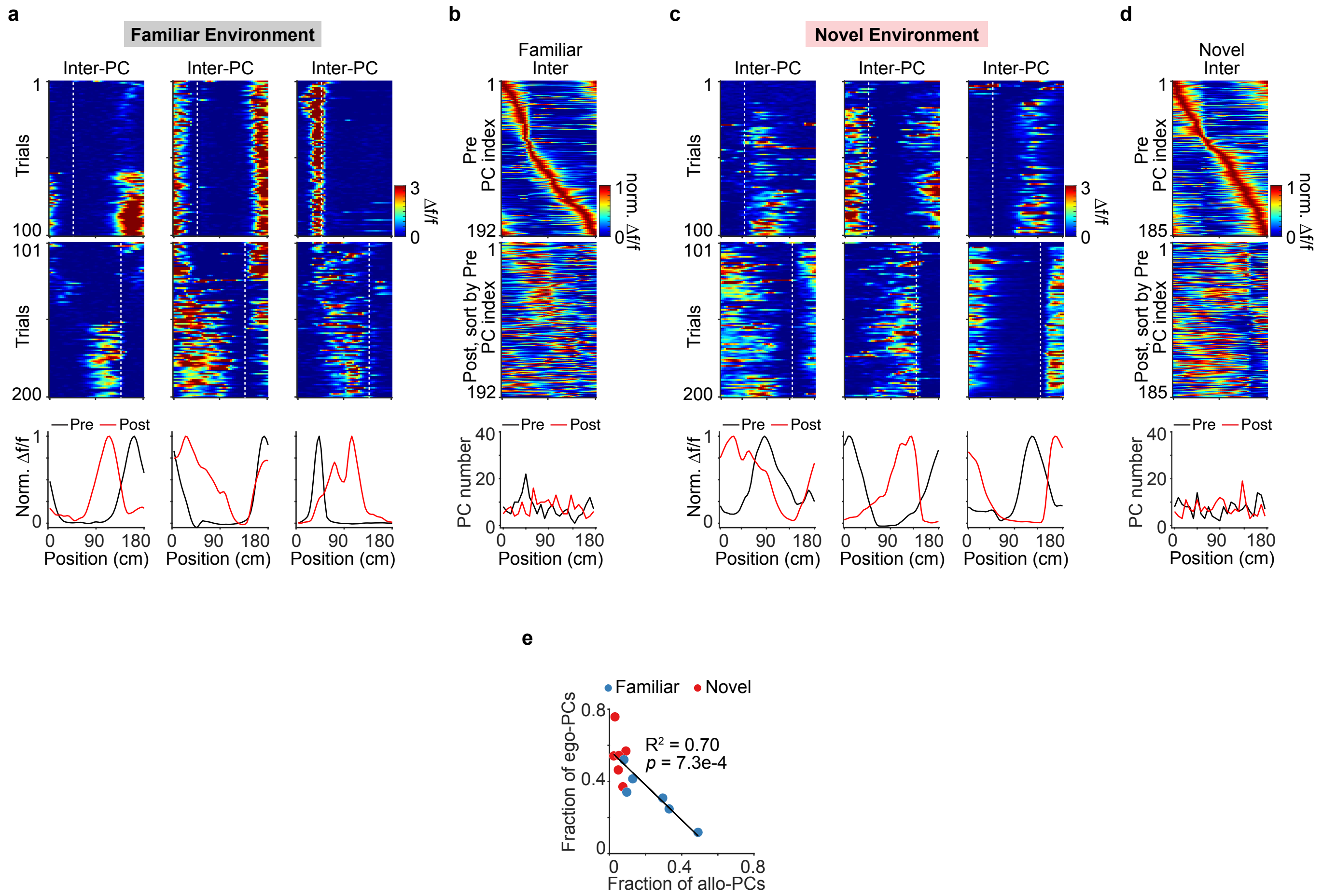


**Extended Data Fig. 3**

**Extended Data Fig. 3 | Intermediate PCs from day 1 on a familiar versus novel belt. a,** Examples of 3 inter-PCs showing their responses to the reward switch. Colormaps show trial by trial Δf/f as a function of position during Pre (top) and Post (middle). White dashed lines mark the reward locations. Bottom: peak-normalized mean Δf/f for each cell during Pre and Post. Pre: before reward switch. Post: after reward switch. **b**, PC representation. Top: colormap of peak-normalized mean Δf/f for all inter-PCs (n = 6 mice) during Pre ranked by PC peak Δf/f location. Middle: colormap of the same inter-PCs using the peak-normalized mean Δf/f from Post, sorted by Pre. Bottom: PC number as a function of position for Pre and Post. **c-d,** Same plots as **a-b** but show 3 examples of inter-PCs on a novel belt on day 1 and the entire PC representation in Pre and Post (n = 6 mice). **e,** The fraction of ego-PCs as a function of the fraction of allo-PCs. Black line is a linear fit, suggesting a strong inverse correlation between the fraction of egocentric and allocentric PCs. n = 12 mice, pooled from both familiar and novel belt groups. Each dot represents one animal.

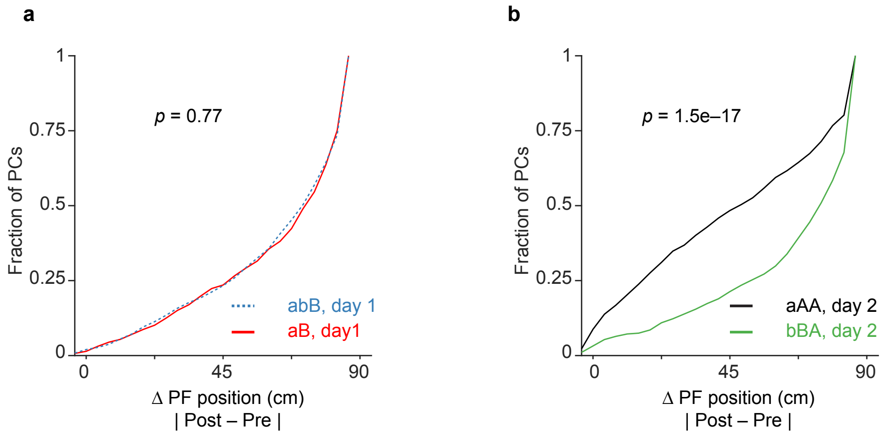


**Extended Data Fig. 4**

**Extended Data Fig. 4 | Experience-dependent, flexible allocentric and egocentric CA1 PC representation. a,** The fraction of PCs as a function of PF position change between the Post and Pre for group 'aB' (solid red, n = 6 mice) and 'abB' (dashed blue, n = 4 mice) on day 1, indicating that the egocentric place representation on a novel belt was not due to lack of experience with fixed reward. 'aB': mice were trained on belt A with fixed reward at 50 cm (lower case 'a'), tested on belt B with reward switch (upper case 'B'). Day 1 for 'aB': the first day of exposure to belt B and the reward switch. 'abB': mice were trained on belt A with fixed reward (lower case 'a'), then trained on belt B with another 4 consecutive days with fixed reward (lower case 'b'), finally tested on belt B with a reward switch (upper case 'B'). Note that 2 of the 4 animals in the 'abB' groups have been exposed to the reward switch on the belt B before the extra 4-day fixed reward training, and the other 2 animals had never experienced the reward switch before the final test. Day 1 refers to the first day of the reward switch on belt B after extensive training experience on belt B with fixed reward. (Two-sample Kolmogorov-Smirnov test, $p = 0.77$). **b,** Same plot as **a** but for group 'aAA' on day 2 (black, n = 6 mice) and 'aBA' on day 2 (green, n = 3 mice), indicating that the balanced spatial coding observed on the familiar belt (Fig.1) was not dictated by the belt cues but was formed with experience. 'aAA': mice were trained on belt A with fixed reward (lower case 'a'), then tested on the familiar belt A for two consecutive days with a reward switch. Day 2 refers to the second day of testing with the reward switch on belt A. 'aBA': mice were trained on the belt B with fixed reward (lower case 'b'), then tested on the familiar belt B with a reward switch (upper case 'B') as day 1, and tested on the belt A (upper case 'A', now novel) on day 2. (Kolmogorov-Smirnov test, $p = 1.5e{-}17$).

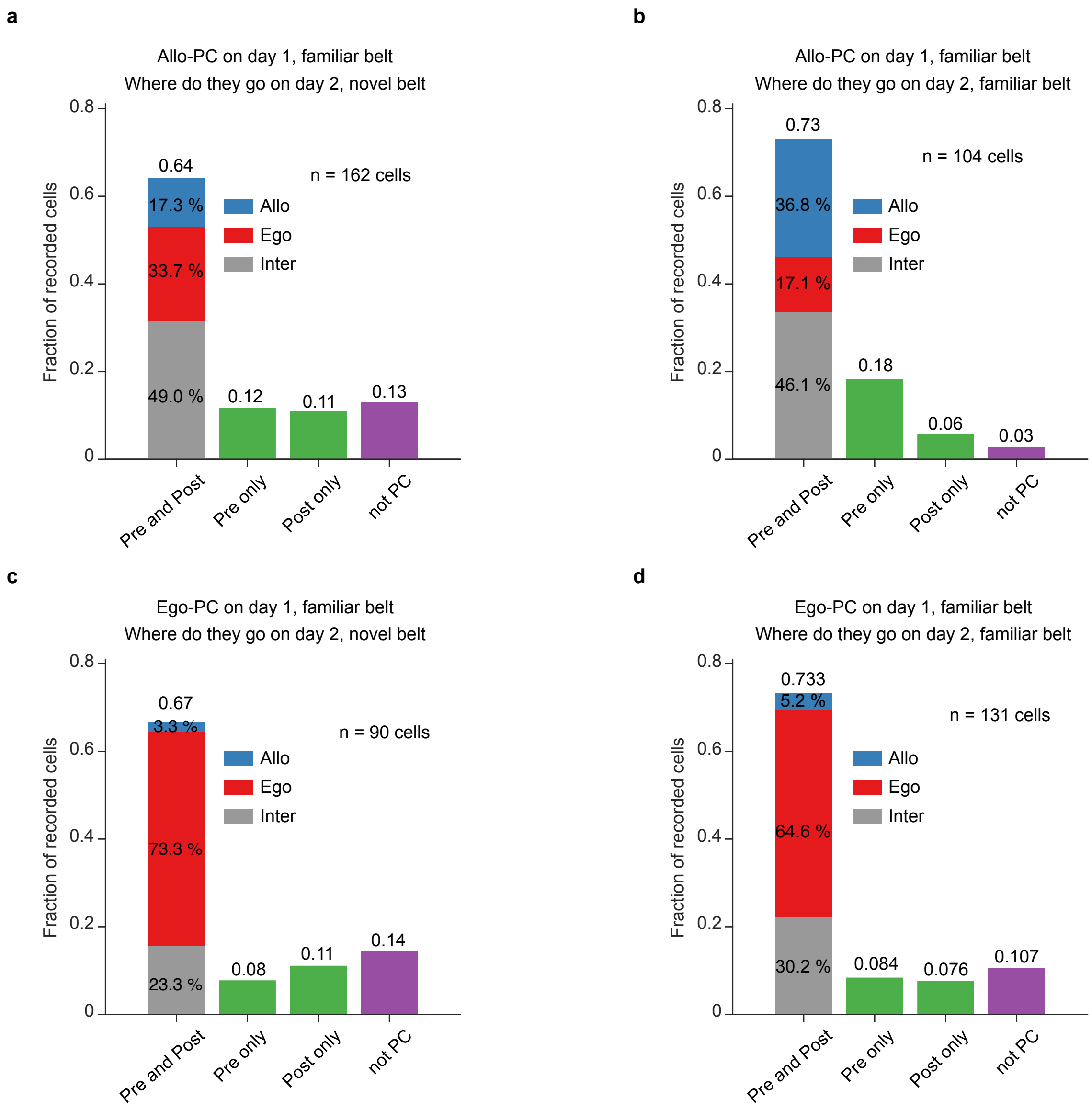


**Extended Data Fig. 5**

**Extended Data Fig. 5 | Longitudinal tracking of cells from day 1 to day 2. a,** Tracking of allocentric PCs across different belts. Bar plots illustrate, of the allocentric PCs identified from day 1 on a familiar belt, the fraction of different types of cells on day 2 on a novel belt (162 allocentric PCs, n = 5 mice). The cells that had reliable PFs during Pre and Post on day 2 were further divided into three categories—allocentric, egocentric and intermediate. Pre: before reward switch. Post: after reward switch. Day 1 and 2 refer to the first and second day of reward switch, respectively. **b,** same plot as **a**, but on the same familiar belt for day 1 and day 2 (104 allocentric PCs, n = 6 mice). **c,** Tracking of egocentric PCs across different belts. Bar plots illustrate, of the egocentric PCs identified from day 1 on a familiar belt, the fraction of different types of cells on day 2 on a novel belt (90 egocentric PCs, n = 5 mice). **d,** same as **c**, but on the same familiar belt for day 1 and day 2 (131 egocentric PCs, n = 6 mice).

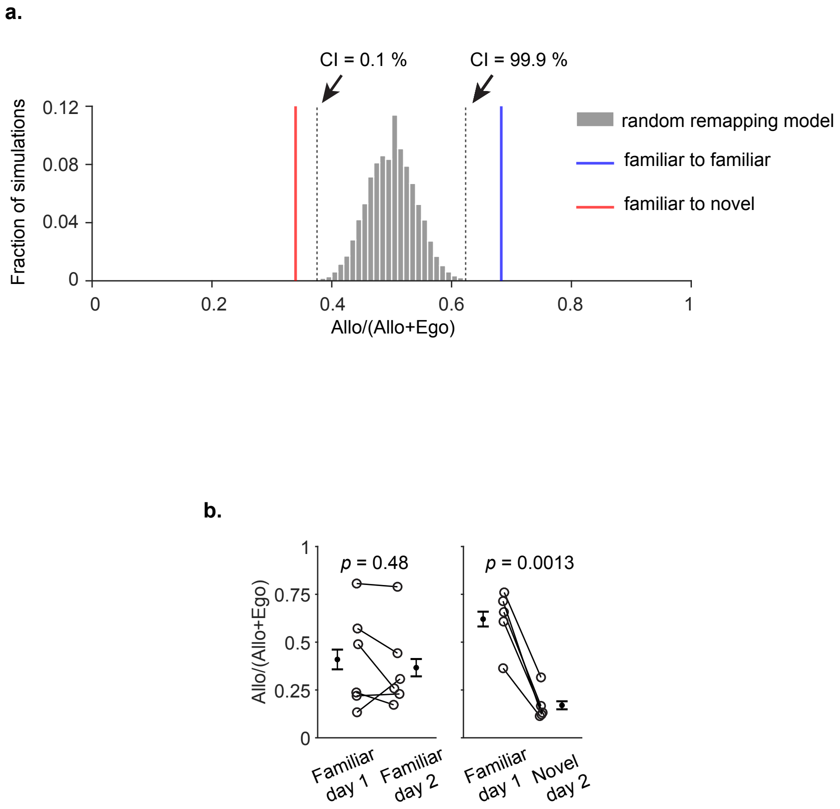


**Extended Data Fig. 6**

**Extended Data Fig. 6 | Stability of allocentric PCs across days. a,** The distribution (grey) describes the fraction of simulations using a random remapping model as a function of an Allo/(Allo+Ego) index. “Allo/(Allo+Ego)” is calculated as the number of allocentric PCs that remained allocentric on next day divided by the total number of allocentric PCs that remained allocentric or became egocentric on next day. Two black vertical dashed lines mark the $0.1^{th}$ and $99.9^{th}$ percentile of the simulations. The blue line marks the observed value for Allo/(Allo+Ego) index when animals experienced the same familiar belt for two days. This value is larger than $99.9^{th}$ percentile of the random remapping population, suggesting that allocentric PCs are significantly more stable than random. The red line marks the observed value for Allo/(Allo+Ego) index when animals switched belts from familiar to novel. This value is smaller than $0.1^{th}$ percentile of the random remapping population, suggesting that allocentric PCs are significantly less stable than random when switching belts. This suggests the fraction of cells that became egocentric from allocentric is significantly higher than expected from a random remapping model. CI: Confidence interval. n = 6 mice. **b,** Comparison of the Allo/(Allo+Ego) index across days for familiar vs familiar belt (left, familiar belt on day 1: 0.41±0.11; familiar belt on day 2: 0.37±0.092; Paired student *t*-test, $p = 0.48$) and familiar to novel (right, familiar belt on day 1: 0.62±0.069; familiar belt on day 2: 0.17±0.038; Paired student *t*-test, $p = 0.0013$). “Allo/(Allo+Ego)” is calculated as the number of allocentric PCs that stayed allocentric on next day divided by the total number of allocentric PCs that either remained allocentric or became egocentric on next day. Each dot represents one animal. The plot showed that Allo/(Allo+Ego) index remained relatively stable within the same environment across two days but decreased significantly from a familiar to a novel environment.

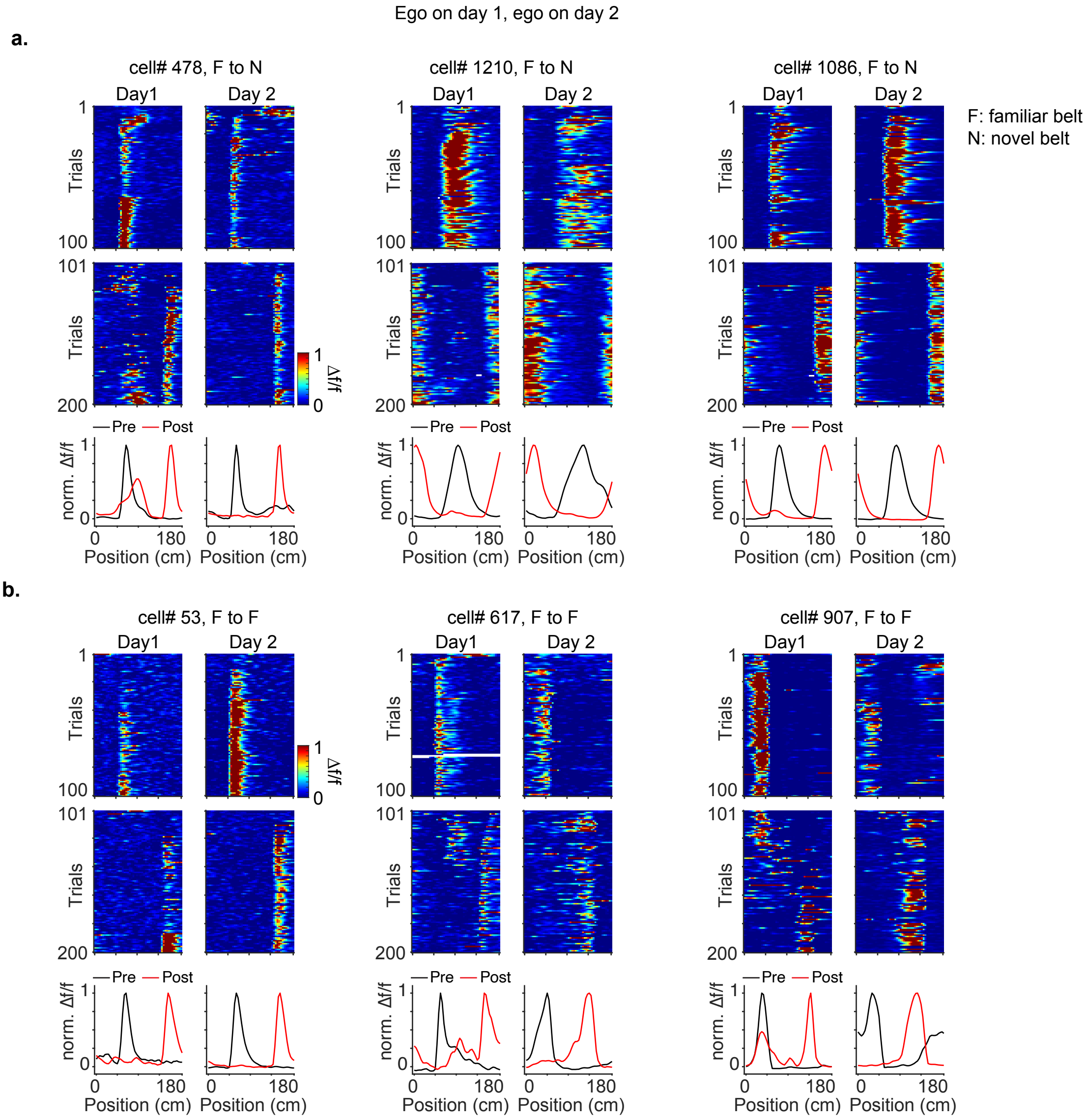


**Extended Data Fig. 7**

**Extended Data Fig. 7 | Stability of egocentric PCs across two days. a,** Three examples of egocentric PCs on a familiar belt on day 1 that remained egocentric on a novel belt on day 2. Colormaps of Δf/f across space before (top, Pre) and after (middle, Post) the reward switch, for day 1 (left) and day 2 (right). Bottom: Peak-normalized (norm.) mean Δf/f across space for Pre and Post, respectively. Day 1 and Day 2: first and second day following reward switch, respectively. **b,** same as **a**, but are examples of egocentric PCs that remained egocentric for two days on the same familiar belt.

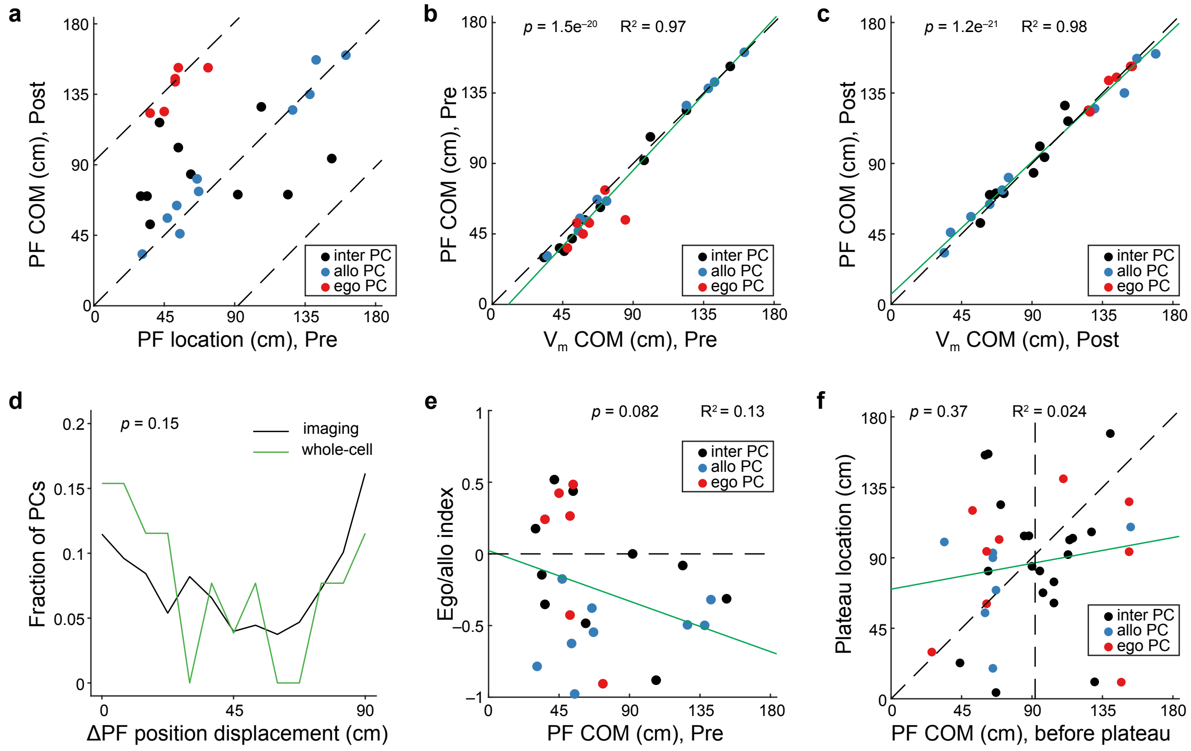


**Extended Data Fig. 8**

**Extended Data Fig. 8 | Other characteristics of CA1 PC remapping. a,** PF remapping of 26 recorded CA1 PCs. The reward is at 184 and 92 cm before (Pre) and after (Post) reward switch, respectively. The dashed lines indicate the distribution center of ego- and allo-centric cells (n = 26 cells). The black, blue and red dots indicate the intermediate, allo- and ego-centric PCs, respectively. **b,** The correlation of $V_m$ COM and the PF COM in Pre. The green line is a linear fit (n = 26 cells, $p = 1.5 \times 10^{-20}$, $R^2 = 0.97$). **c**, The correlation of $V_m$ COM and the PF COM in Post. The green line is a linear fit. (n = 26 cells, $p = 1.2 \times 10^{-21}$, $R^2 = 0.98$). **d**, The fraction of PCs as a function of shifted PF distance after reward switch. The black and green lines are from imaging and whole-cell data, respectively ($p = 0.15$, two-sample Kolmogorov-Smirnov test). **e**, The ego/allo index as a function of PF COM in Pre. The reward is at 184 cm in Pre. The green line is a linear fit (n = 25 cells, $p = 0.08$, $R^2 = 0.13$). **f**, The correlation of PF COM before spontaneous plateaus and the location of spontaneous plateaus. The green line is a linear fit (n = 35 plateaus, $p = 0.37$, $R^2 = 0.024$).

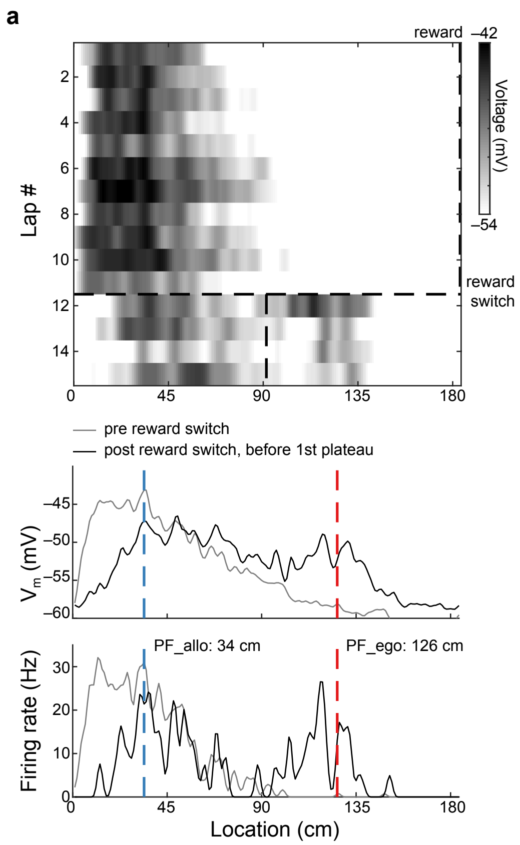


**Extended Data Fig. 9**

**Extended Data Fig. 9 | An example cell demonstrating the split of a place field into ego- and allocentric $V_m$ ramps immediately following reward switch. a,** Top: the shaded heatmap shows the $V_m$ in space across trials before (Pre) and after (Post) reward switch. The dashed lines indicate the reward locations (at 184 and 92 cm in Pre and Post, respectively). Middle: the averaged $V_m$ of Pre (grey) and Post (black) trials. The blue and red dashed lines indicate original (allocentric) PF and the egocentric PF (92 cm away from original PF), respectively. Bottom: the mean spatial firing rate of the cell of Pre (grey) and Post (black) trials.

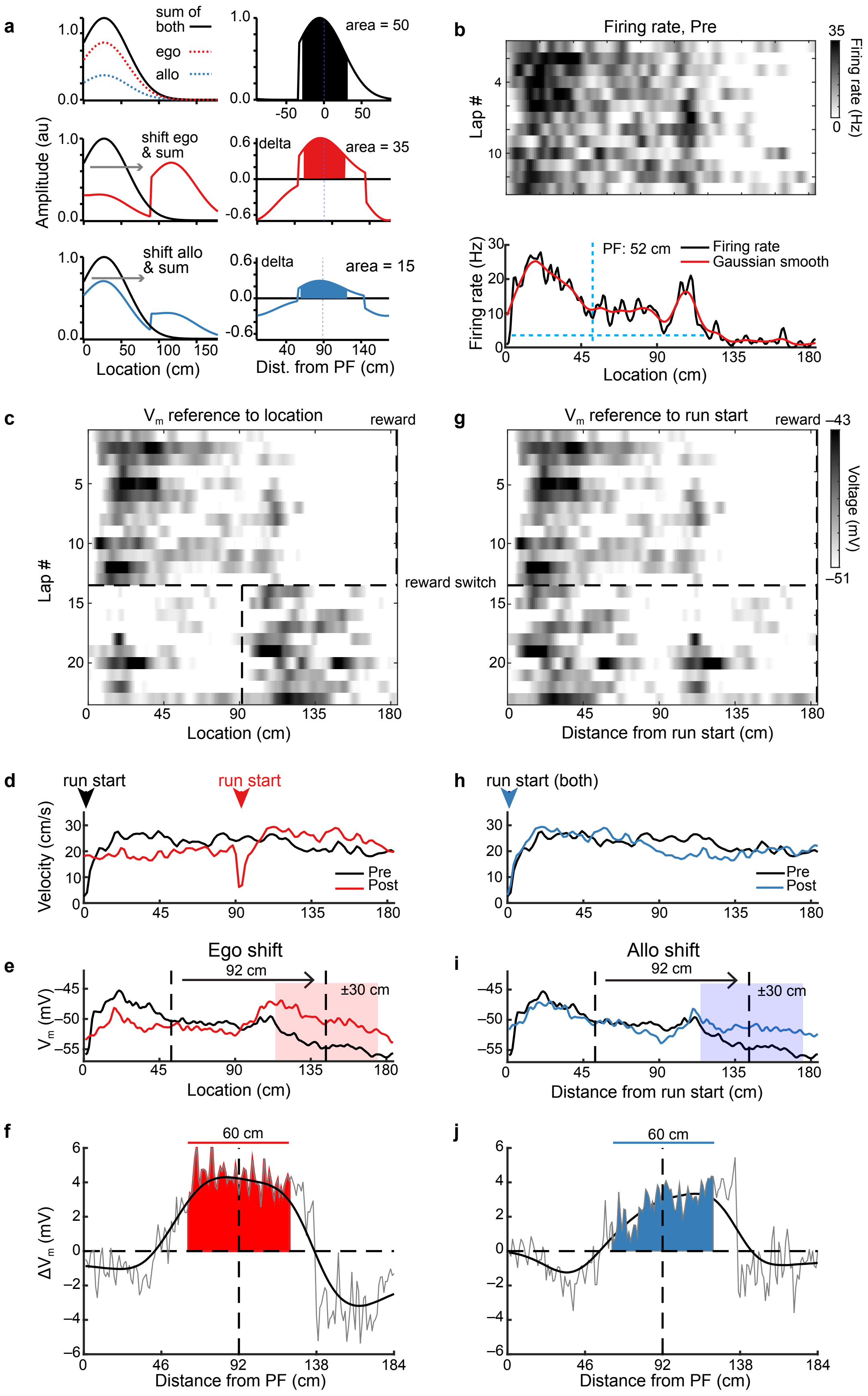


**Extended Data Fig. 10**

**Extended Data Fig. 10 | Calculating the putative ego- and allo-centric $V_m$ by subtraction. a**, A simple illustration of how the ego $V_m$ and allo $V_m$ was calculated. The top shows a PF (left, black) comprised of two components: ego $V_m$ (left, red) and allo $V_m$ (left, blue), and the total area of the PF is 50 (right, aligned to the PF location). The middle left shows the summation of $V_m$ (red) when shifting the ego $V_m$, thus revealing the putative ego $V_m$ (red, shifted $V_m$) with an area of 35 from the subtraction of red and black $V_m$ traces in the left (right). Similarly, the bottom left shows the summation of $V_m$ (blue) when shifting the allo $V_m$, revealing the putative allo $V_m$ (blue, shifted $V_m$) with an area of 15 from the subtraction of blue and black $V_m$ traces in the left (right). **b,** The firing rate of an example PC before reward switch (Pre, reward at 184 cm). The top shaded heatmap shows the firing rate across trials before reward switch. The bottom is the averaged firing rate (black), and the red line indicates the Gaussian smoothed representation of the averaged firing rate. The blue dashed horizontal line indicates the extent of the PF. The blue dashed vertical line marks the COM of the PF, which defines PF location. **c**, A shaded heatmap shows the $V_m$ in space of the same cell in **b** across trials before (Pre) and after (Post) reward switch, referenced to physical locations of the track. Two dashed vertical lines indicate the reward locations (at 184 and 92 cm in Pre and Post, respectively). The dashed horizontal line marks the trial of reward switch. **d**, The running trajectory in Pre (black) and Post (red). Note the run start shifted ~92 cm (from 0 to 92 cm) after the reward switch. **e**, The averaged $V_m$ traces of Pre (black) and Post (red) trials from **c**. The dashed lines indicate that the COM of ego $V_m$ shifted 92 cm (now at 144 cm) from the original PF (52 cm), due to the shift of the run start in **d**. The red shading (±30 cm around the COM of shifted ego $V_m$) marks the area of ego $V_m$. **f**, The subtraction of the $V_m$ traces from **e**. 0 cm marks the COM of original PF in Pre. The grey and black lines are the raw and the Gaussian smoothed traces, respectively. The red shading area (±30 cm around the COM of shifted ego $V_m$) quantifies the magnitude of ego $V_m$. **g–j**, Same as **c–f**, but the $V_m$ traces are referenced to the start of running. Now the reward is at 184 cm during Pre and Post in **g**, and the running start location is at 0 cm during Pre and Post in **h**. Note the $V_m$ in Post in **g** shifted 92 cm compared to **c**. Thus, the $V_m$_allo shifted 92 cm due to the alignment to the start of running in **i**. The blue shading area in **j** (±30 cm around the COM of shifted allo $V_m$ subtracted from **i**) quantifies the magnitude of allo $V_m$.

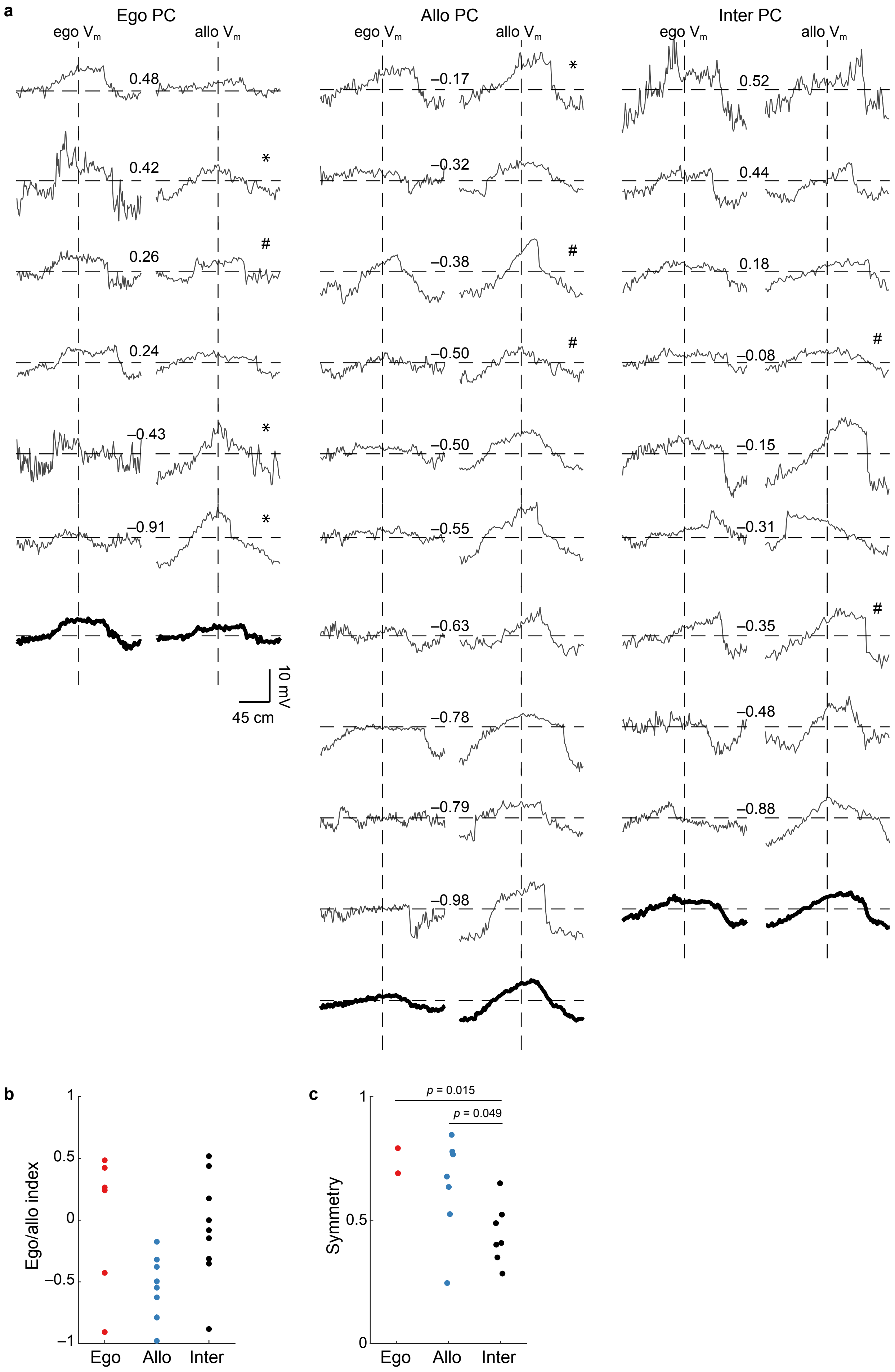


**Extended Data Fig. 11**

**Extended Data Fig. 11 | Putative ego- and allo-centric $V_m$ in individual CA1 PCs. a,** The ego- and allo-centric $V_m$ in 25 recorded CA1 PCs (one cell is excluded due to the immediate spontaneous plateau after reward switch, see Methods), grouped by PC category. The number indicates the ego/allo index of a given cell. The bold traces at the bottom are the averaged $V_m$ for each individual column (cells marked by the star are excluded). The star marks the cells that changed their categories due to spontaneous plateau potentials. The # symbols mark the cells with two separate PFs (see Methods). **b**, The ego/allo index in ego-, allo-centric and intermediate cells. **c**, The symmetry of the total $V_m$ (ego $V_m$ + allo $V_m$) in ego-, allo-centric and intermediate cells (the 9 cells marked by the star and # symbol in **a** are excluded. 0.74±0.051, 0.64±0.77 and 0.44±0.46 for ego-, allo- centric and intermediate cells, respectively, $p$ = 0.015 and 0.049 for ego vs. inter and allo vs. inter cells, unpaired $t$-test).

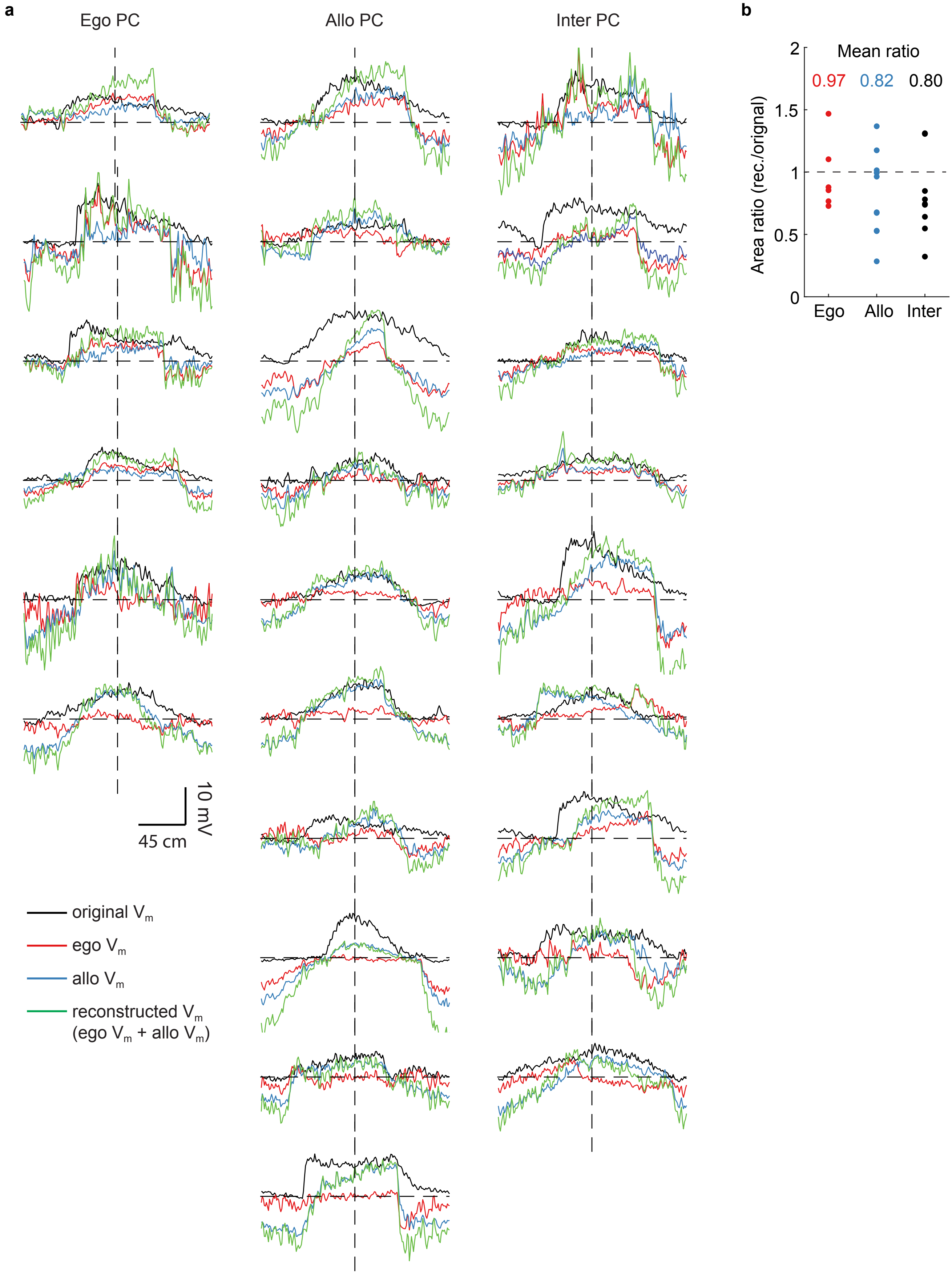


**Extended Data Fig. 12**

**Extended Data Fig. 12 | The reconstructed PF $V_m$ and original PF $V_m$ in individual CA1 PCs. a,** The original, egocentric, allocentric and reconstructed $V_m$ (ego $V_m$ + allo $V_m$) in 25 recorded CA1 PCs (one cell is excluded due to an immediate spontaneous plateau after reward switch, see Methods), grouped by PC category. **b**, The area ratio of the reconstructed PF $V_m$ versus the original PF $V_m$ in ego, allo and inter PCs. The mean ratios are 0.97, 0.82 and 0.80 for ego-, allo- centric and intermediate cells, respectively.

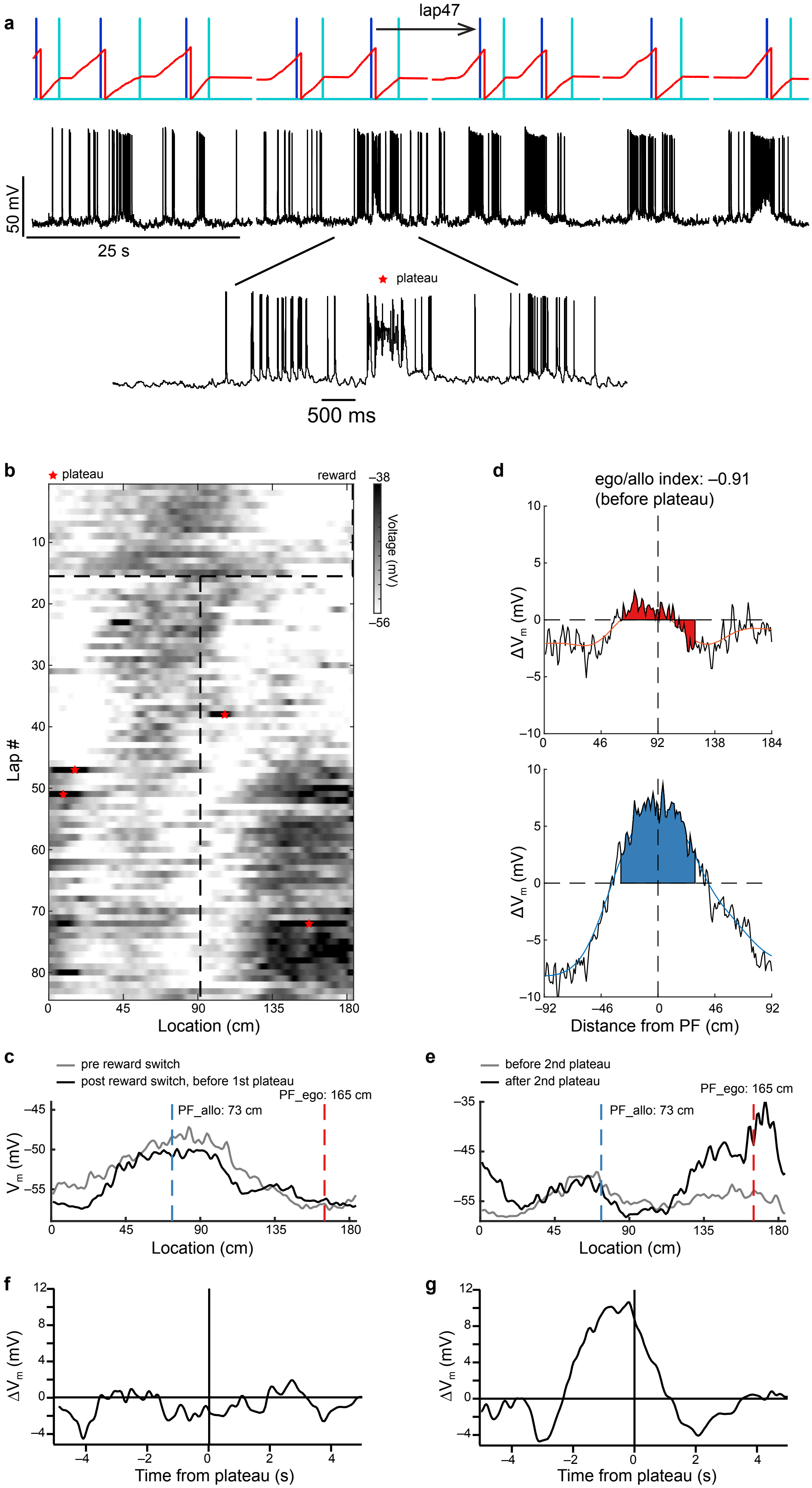


**Extended Data Fig. 13**

**Extended Data Fig. 13 | Spontaneous plateau potential initiation after reward switch. a,** $V_m$ of a PC with spontaneous plateau potentials (black). The red line indicates the location of the mouse on the belt (from 0 to 184 cm). The light blue lines indicate the reward location (at 92 cm after reward switch (Post)) and the dark blue lines indicate the end of the lap (184 cm). Bottom: $V_m$ trace expanded showing the spontaneous plateau potential at the beginning of lap 47. **b,** A shaded heatmap shows the $V_m$ across trials before (Pre) and after (Post) reward switch. The vertical dashed lines indicate the reward locations (at 184 and 92 cm in Pre and Post, respectively). The horizontal dashed line marks the trial of the reward switch. The red stars indicate the spontaneous plateau potentials. **c**, The averaged $V_m$ traces of Pre (grey, lap 1–15) and Post (black, before 1st plateau, lap 16–37) trials. The blue and red dashed lines indicate the original (allocentric) PF and the egocentric PF (92 cm away from original PF). **d**, The putatively ego- (top, red) and allocentric (bottom, blue) $V_m$ of this example cell. **e**, The averaged $V_m$ traces before (grey, trials 39–46) and after (black, lap 48–50) the 2nd plateau potentials (at lap 47). The blue and red dashed lines indicate the original (allocentric) PF and the egocentric PF (92 cm away from original PF). **f**-**g**, The BTSP kernels of 1st (**f**, lap 38) and 2nd (**g**, lap 47) plateau potentials.
